# Behavioral investigation of allocentric and egocentric cognitive maps in human spatial memory

**DOI:** 10.1101/2025.01.17.633375

**Authors:** Laura Nett, Tim A. Guth, Philipp K. Büchel, Nuttida Rungratsameetaweemana, Lukas Kunz

## Abstract

Spatial memory is a fundamental cognitive function that enables humans and other species to encode and recall the locations of items in their environments. Humans employ diverse strategies to support spatial memory, including the use of cognitive maps. Cognitive maps are mental representations of the environment that organize its content along two or more continuous dimensions. In allocentric cognitive maps, these dimensions form a Cartesian coordinate system referenced to the environment. In egocentric cognitive maps, the dimensions form a polar coordinate system centered on the subject. To better understand how humans employ allocentric and egocentric cognitive maps for spatial memory, we performed a behavioral study with a novel task designed to directly and explicitly assess both types of cognitive maps. During encoding periods, participants navigated through a virtual environment and encountered objects at different locations. During recall periods, participants aimed at remembering these locations in abstract allocentric and egocentric coordinate systems. Our results show that relationships between the objects and the environment, such as their distance to boundaries and corners, were associated with allocentric memory performance. Relationships between the objects and the participant, including their distance and orientation to the participant’s starting position, were linked to egocentric memory performance. Spatial feedback during recall supported performance within allocentric and egocentric domains, but not across domains. These findings are compatible with the notion that allocentric and egocentric cognitive maps operate as (partially) independent systems for spatial memory, each specialized in processing specific types of spatial relationships.

## 1. Introduction

Spatial memory enables humans and other species to remember the locations of objects within their surroundings, making it a fundamental cognitive function for daily life (Allen, 2013; Ekstrom et al., 2018; Ekstrom and Ranganath, 2018). To achieve effective spatial memory, humans often rely on cognitive maps that provide more or less coherent and comprehensive mental representations of the environment (Burgess, 2006). Based on a wide range of behavioral, electrophysiological, neuroimaging, and lesion studies (e.g., Tolman, 1948; O’Keefe, 1991; Kitchin, 1994; Nadel, 2013; Epstein et al., 2017; Grieves and Jeffery, 2017; Behrens et al., 2018; Bellmund et al., 2018; Bottini and Doeller, 2020; Peer et al., 2021; Jeffery, 2024; Jeffery et al., 2024; Sharp, 2025), cognitive maps are thought to manifest in two distinct forms by representing the environment in allocentric or ego-centric reference frames, although most studies have used the concept of cognitive maps to refer only to allocentric cognitive maps (Tolman, 1948; Epstein et al., 2017). These allocentric cognitive maps are world-referenced and thus store information about locations and directions in a Cartesian coordinate system anchored to the external environment, with the subject moving through this map. For instance, a person might recall that their home is located in the western part of their hometown. In contrast, in egocentric cognitive maps, the spatial dimensions form a polar coordinate system that is centered on the subject, representing spatial information in relation to the person’s current position and orientation. As the subject moves through an environment, the egocentric coordinate system remains anchored to their vantage point, causing the environment to shift relative to it. For example, a person might recall that their home is two kilometers to the right of their current location and orientation. After turning around, it is represented as two kilometers to the left. Together, these two types of spatial representations help humans recall spatial memories in various situations. However, the exact relationship between allocentric and egocentric cognitive maps and their interactions during spatial memory processes remain unclear.

The distinction between allocentric and egocentric cognitive maps is supported by a large number of neurophysiological studies that identified specialized cell types representing various types of spatial information (Grieves and Jeffery, 2017; Bicanski and Burgess, 2020). Place cells, grid cells, and head-direction cells (O’Keefe and Dostrovsky, 1971; Taube et al., 1990; Hafting et al., 2005) are among the most important cell types that are believed to underlie allocentric cognitive maps as they encode a subject’s locations and directions relative to the environment (McNaughton et al., 2006). For example, a place cell may show elevated spiking activity whenever an animal is in the east part of the environment. Place, grid, and head-direction cells have been studied most often in freely moving rodents (Moser et al., 2017), with some evidence for similarly tuned neurons in humans (Jacobs et al., 2013; Miller et al., 2013; Qasim et al., 2019, 2021; Kunz et al., 2024). Neurons that may be involved in the neural basis of egocentric cognitive maps comprise cells that increase their firing rates whenever a boundary, a landmark, or some other reference point is at a particular egocentric direction and distance from the subject (Bicanski and Burgess, 2020; Kunz et al., 2021). This includes recently discovered egocentric boundary vector cells (Wang et al., 2018; Hinman et al., 2019; Alexander et al., 2020), egocentric center-bearing cells (LaChance et al., 2019; LaChance and Taube, 2023), and egocentric item-bearing cells (Wang et al., 2018). Again, a few studies have obtained evidence for similarly tuned neurons in the human brain (Jacobs et al., 2010; Kunz et al., 2021).

To what extent allocentric and egocentric cell types are distinct from each other, how much they overlap regarding their anatomical distribution, and how they communicate with each other are largely unresolved questions. A common framework is that the hippocampus and surrounding medial temporal lobe structures play a key role in allocentric processing, while egocentric processing is more closely associated with the parietal cortex (Byrne et al., 2007; Bicanski and Burgess, 2018; Wang et al., 2020). Direct evidence for this framework is difficult to obtain, however, and it remains elusive to what extent the different cell types are involved in either allocentric or egocentric cognitive maps or both. Among others, this is because allocentric and egocentric processes are difficult to discern at the behavioral level (Burgess, 2006; Ekstrom and Hill, 2023). Behavioral tasks testing the role of allocentric and egocentric cognitive maps in spatial memory as independently as possible may thus help delineate allocentric and egocentric neural processing (Krakauer et al., 2017).

Previous research developed a wide range of tasks to assess allocentric and egocentric spatial memory in a variety of species, such as the Morris water maze (Morris, 1981; Vorhees and Williams, 2006, 2014; Jacobs et al., 2016). A useful way of testing allocentric memory in humans is map drawing (Ekstrom and Isham, 2017). After exploring an environment from a first-person perspective, participants are tasked with drawing a map of the environment from a bird’s eye view (Jafarpour and Spiers, 2017; Chiang et al., 2023). This task quantifies the extent to which participants are able to integrate individual, discontinuous experiences into a coherent framework that is organized by continuous dimensions (Bellmund et al., 2018; Behrens et al., 2018). Although the act of drawing the allocentric map may involve interconnecting egocentric viewpoints, its outcome is explicitly allocentric and requires a spatial representation that is independent of the observer’s current location or orientation, focusing on the relationships between objects in the environment relative to one another (Ekstrom and Isham, 2017). Drawing a map of an environment or localizing objects within such a map may thus constitute one of the most direct behavioral readouts of someone’s allocentric understanding of an environment.

Many studies also developed tasks specialized in testing egocentric spatial memory (Wang and Spelke, 2002; Burgess, 2006; Filimon, 2015). For example, in the scene and orientation-dependent pointing task, participants experience an environment from a first-person perspective and are asked to point in the direction of goal locations that are out of view (Mou et al., 2004; Zhang et al., 2014; Ekstrom and Isham, 2017). This and similar tasks require abstract spatial knowledge of the environment anchored to specific locations and orientations of the individual, which may preferentially form during important behavioral events (e.g., upon entering a new environment). Asking individuals to localize objects within abstract maps organized by the dimensions of distance and direction relative to an individuals remembered first-person perspective may thus enable direct readouts of egocentric cognitive maps. Such egocentric spatial knowledge is often used in everyday situations and may be particularly useful when there is insufficient experience to construct an allocentric cognitive map (Burgess, 2006; Zhang et al., 2014). Although egocentric relationships between a person and objects in the environment can, in principle, be derived from allocentric knowledge of an environment (Ekstrom et al., 2014), previous research indicates that egocentric cognitive maps are distinct from and coexist with allocentric cognitive maps (Ekstrom et al., 2014; Ekstrom and Isham, 2017).

Here, we developed a behavioral task, the “Garden Game” (Figures 1 and 2), as a new resource to differentiate spatial memory based on allocentric and egocentric cognitive maps and to study how humans use these maps during spatial memory recall. The Garden Game is a laptop-based virtual-reality task, which is freely available and modifiable (Materials & Methods). In each trial during encoding, participants navigate through a virtual environment (a garden) and aim at memorizing the locations of two objects (animals) within the environment. During subsequent retrieval, participants are presented with two abstract coordinate systems to recall the object locations relative to the external environment (allocentrically) and relative to their initial starting position (egocentrically). We suggest that this design provides a means to quantify the use of allocentric and egocentric cognitive maps in spatial memory. Additionally, the Garden Game enables the investigation of various behavioral, environmental, item-related, and demographic determinants of spatial memory. We propose that by demonstrating similar versus different effects of these determinants on allocentric versus egocentric spatial memory, the Garden Game helps understand to what extent allocentric and egocentric cognitive maps are overlapping or distinct entities.

**Figure 1:**
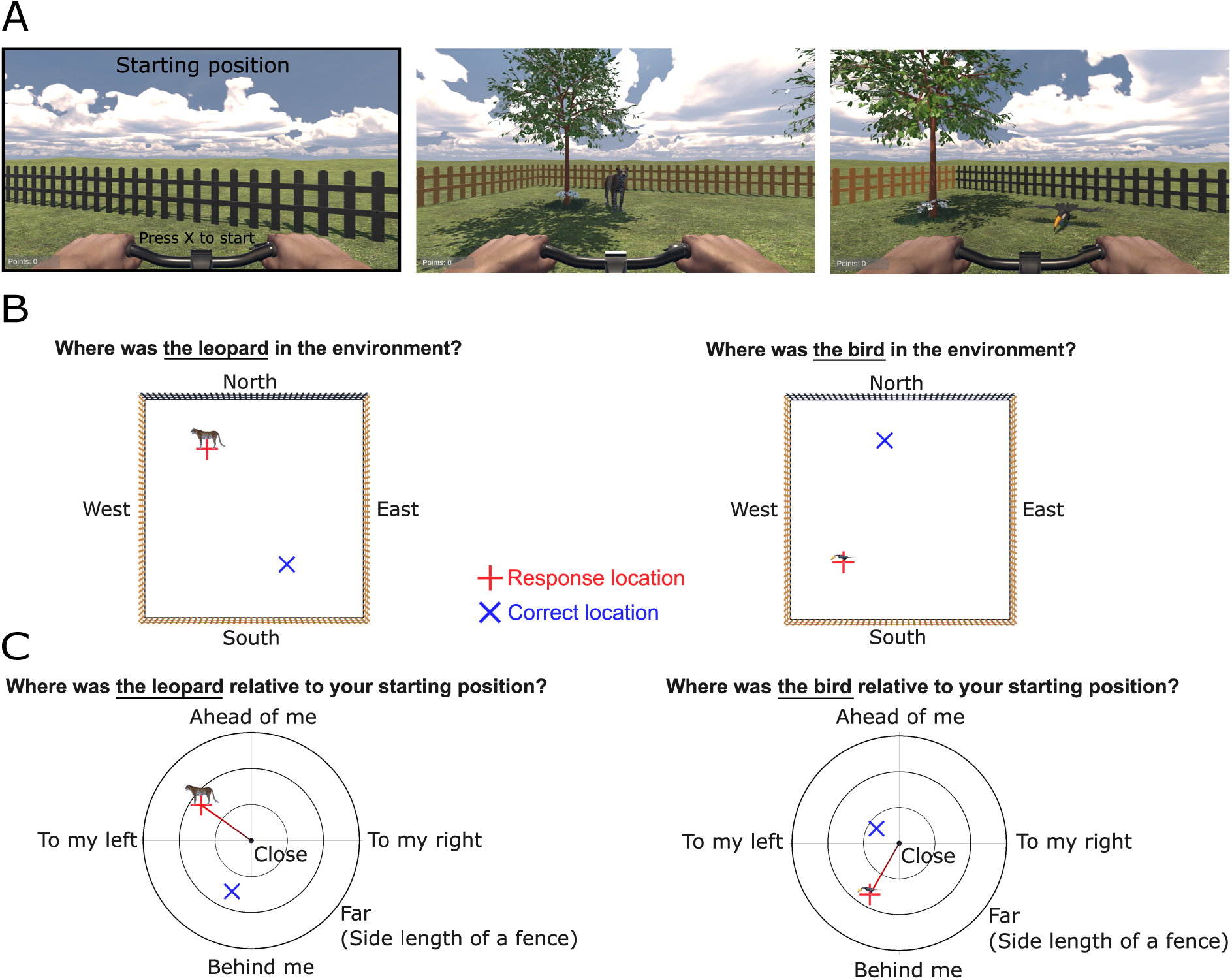
Garden Game task. (A) During the encoding period of each trial, participants freely navigate a virtual garden environment and encounter two objects (animals) at different locations within the environment. Participants are instructed to encode the object locations relative to the external environment (allocentrically) and relative to their initial starting position (egocentrically). (B–C) During the retrieval period of each trial, participants are asked to recall the locations of the objects from encoding in abstract allocentric and egocentric coordinate systems. The order of allocentric and egocentric recall is counterbalanced across trials. (B) During allocentric retrieval, participants recall the positions of the two objects relative to the environment. Participants move a red cross across the screen and press a button when they have reached the remembered location of the object. Immediately following the response, participants receive spatial feedback in the form of a blue cross indicating the correct location of the object, as shown in the figure. (C) During egocentric retrieval, participants recall the positions of the two objects relative to their starting position from encoding. Again, they move a red cross across the screen and press a button when they have reached the remembered location of the object. Immediately following the response, participants receive spatial feedback in the form of a blue cross, which indicates the correct location of the object, as shown in the figure.

**Figure 2:**
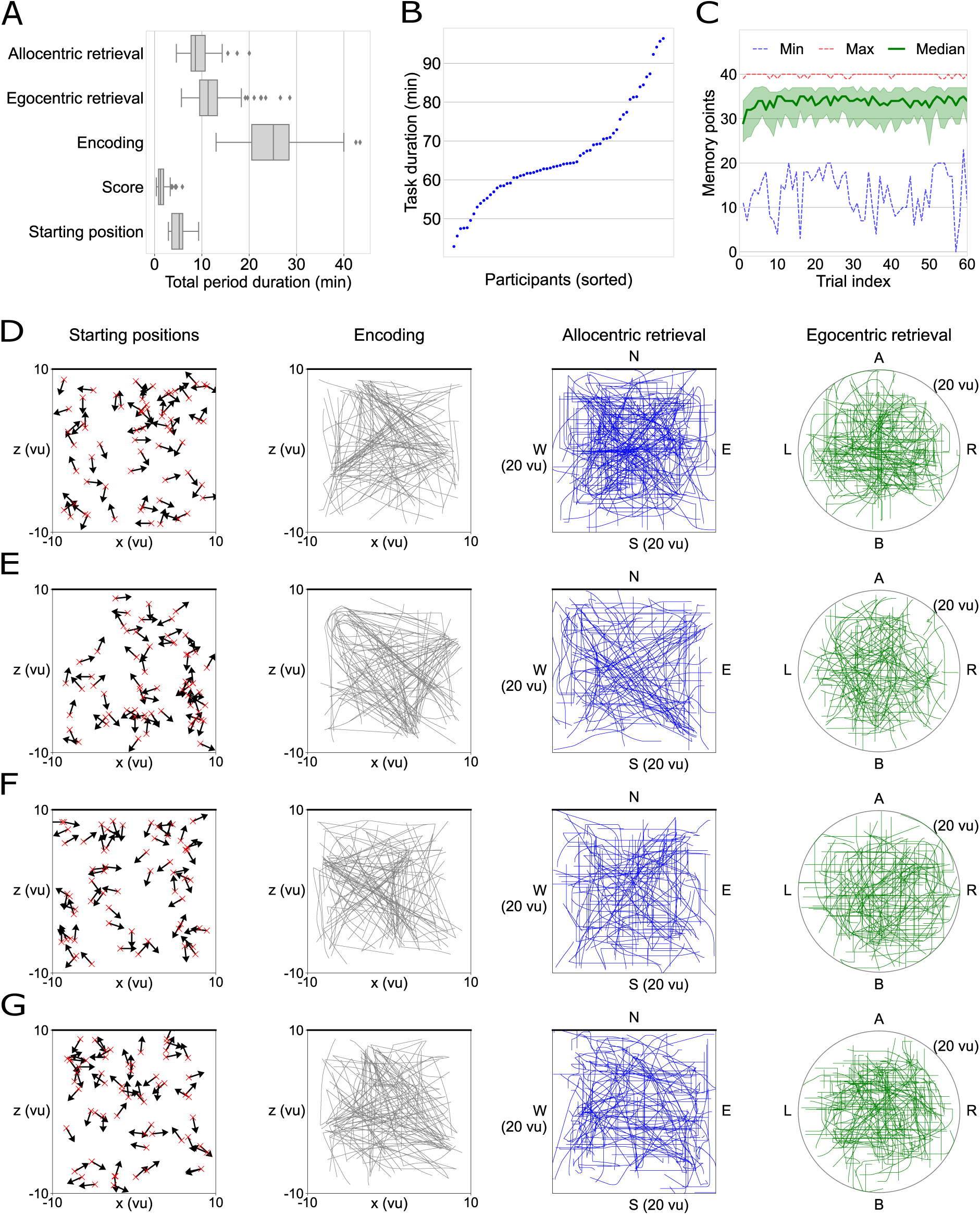
Basic behavioral results. (A) Total duration of each trial period. Boxplots show the median across participants as thick black line, the interquartile range as box, minimum and maximum as whiskers, and outliers as dots. (B) Overall task duration. Each dot represents one participant. (C) Total memory points per trial. Thick green line, median; green shaded area, median ± interquartile range; red dotted line, maximum; blue dotted line, minimum. (D–G) Example behavior of four randomly selected participants. First column, allocentric starting positions (red crosses) and orientations (black arrows) for all trials. Second column, navigation trajectory during all encoding periods (coverage is largely determined by the starting positions, the locations of the objects, and the number of trials). Third column, response cursor trajectory during all allocentric retrieval periods. Fourth column, response cursor trajectory during all egocentric retrieval periods. A, ahead; B, behind; L, left; R, right. N, north; E, east; S, south; W, west. vu, virtual units.

To foreshadow our findings, we observed that recall performance improved for both egocentric and allocentric memory over the course of the task. Allocentric memory was better for objects close to the boundary and the corners, while egocentric recall was better for objects close to and ahead of the participant’s starting position. Feedback during allocentric retrieval improved subsequent allocentric retrieval, and egocentric feedback stabilized egocentric retrieval. Allocentric feedback did not support egocentric retrieval and *vice versa*. Time elapsed between encoding and retrieval was associated with poorer performance. In a subset of participants where we collected eye-tracking data, viewing the environmental boundaries and corners was associated with better allocentric retrieval, whereas egocentric memory was positively correlated with a longer viewing time of the objects. Across participants, memory performance decreased with older age, and men showed better performance on egocentric recall compared to women. We suggest that the Garden Game provides a measure to distinguish allocentric and egocentric cognitive maps behaviorally, and it may provide a useful resource for studying the neural basis of these different types of cognitive maps (Krakauer et al., 2017).

## 2. Materials & Methods

### 2.1. Ethics approval

All procedures were performed in compliance with relevant laws and institutional guidelines and were approved by the local ethics committee of the University Hospital Bonn (reference number, 261/23-EP; date of approval, September 27, 2023). The privacy rights of human participants were observed and written informed consent was obtained from all participants.

### 2.2. Task

The Garden Game task is a laptop-based virtual-reality task programmed in Unity (Unity Technologies, San Francisco, CA, USA), whose source and build versions are available for download from GitHub (https://github.com/BonnSpatialMemoryLab/ GardenGameTask) and Zenodo. The Garden Game version of this study contained 60 trials, with an additional training trial with verbal instructions at the beginning that was excluded from all analyses. Each trial consists of two parts, encoding and retrieval.

During the encoding periods, the Garden Game features a virtual environment that extends into infinity (no distal cues). The navigable area of the environment is limited by a square boundary with an edge length of 20 virtual units (vu). Virtual units roughly correspond to virtual meters given the height of the player and its movement speed. The square boundary consists of four fences at each cardinal direction. The north boundary (fence) is highlighted in a different color (black) than the other three boundaries (brown), thus serving as an allocentric direction signal. The minimum distance between the participant’s position and a boundary is 0.1 vu. Three trees serve as intramaze landmarks and are placed at the centers of three quarters of the environment (randomized across participants).

At the beginning of each encoding period, participants are located at a random starting position (x and z coordinates ranging from -9 to 9 vu) within the environment and face a specific allocentric direction (minimum duration of 3 seconds; for examples, see the left column of Figure 2, D–G). The starting position is at least 5 vu away from both objects and at least 1 vu away from the intramaze landmarks (trees). Participants are instructed to pay attention to this starting position as they are required to perform ego-centric retrieval relative to this starting position. Next, they start navigating through the virtual garden environment and encounter, in sequence, two objects (animals) at different locations within the environment (x and z coordinates ranging from -8.5 to 8.5 vu; Figure 1A). When reaching an object (cutoff distance of 1 vu), participants remain at this virtual position for a duration of 2 seconds, during which the name of the object is displayed (e.g., “Bird”) and its sound is played. The object then disappears and the next object appears at a different location in the environment. Participants are instructed to memorize the positions of the two objects relative to the participants’ starting position and orientation (egocentrically) and relative to the environment (allocentrically). Participants move through the environment using a gamepad that allows them to move ahead, turn left, and turn right.

During the retrieval periods, the Garden Game displays two abstract coordinate systems for allocentric and egocentric retrieval (Figure 1, B and C). There are four individual recalls per retrieval period (one allocentric recall for each object and one egocentric recall for each object). During allocentric retrieval, participants view an allocentric map of the garden environment from a bird’s eye view (the black north boundary is shown at the top) and a question at the top of the screen indicates which object the participant is asked to retrieve. Participants then move a red cross across the allocentric map and press a button to indicate the remembered allocentric location of the object. Once participants have pressed the response button, they receive spatial feedback in the form of a blue cross appearing at the correct location on the allocentric map (duration of 2 seconds). Additionally, participants’ total score, displayed in the lower left corner, is updated (0–10 points per retrieval) depending on the accuracy of their response. During egocentric retrieval, participants view an egocentric map (i.e., a polar coordinate system with its angles representing egocentric directions and its radii representing egocentric distances) that is centered on their starting position and orientation (ahead is shown at the top). A question at the top of the screen indicates which object the participant is asked to retrieve. Participants then move a red cross across the egocentric map and press a button to indicate the remembered egocentric location of the object relative to their starting position and orientation. Once they have pressed the response button, a blue cross appears in the correct location on the egocentric map to provide spatial feedback, and participants’ total score is updated (0–10 points per retrieval) depending on the accuracy of their response.

For both types of retrieval, memory performance is calculated based on the Euclidean distance between the participant’s response location and the correct location of the object in the abstract reference frames. This Euclidean distance is then ranked within 1 million surrogate distances to account for the fact that different object locations have different chance levels, following established procedures (Miller et al., 2018; Kunz et al., 2021, 2024). A memory-performance rank of 1 represents perfect memory performance, while a rank of 0 represents the poorest memory performance. To keep participants motivated during the task, participants receive points for each response that are added to their total score. The number of points depends on the response-specific memory performance (minimum of 0 points; maximum of 10 points). Points are given based on an adaptive algorithm so that participants generally receive between 7 and 8 points per retrieval (Figure 2C). This algorithm calculates the mean allocentric/egocentric memory performance throughout the task and updates an allocentric/egocentric adjustment value after each trial. The adjustment values are initially set to zero and undergo incremental changes of ±1 based on the mean allocentric/egocentric memory performance. If the mean performance is below 7, the adjustment value becomes more positive, while it becomes more negative if the mean performance is above 8. If the mean memory performance falls between 7 and 8, the adjustment score remains unchanged. Each retrieval type (allocentric and egocentric) has its own adjustment value, ranging from -10 (indicating good performance) to +10 (indicating poor performance).

The order of allocentric and egocentric retrieval alternates between trials and is randomized and counterbalanced across participants. The order of retrieval for the firstly and secondly encoded object is random across all retrieval periods. For each participant, the Garden Game features six different objects (animals), randomly chosen from a pool of 14 possible objects (bird, camel, cat, chicken, dog, elephant, horse, leopard, penguin, pig, pug, rhino, sheep, and tiger). Across all 60 trials of the task, participants encode each object 20 times and recall each object 40 times (20 times allocentrically and 20 times egocentrically). Three of the objects have the same allocentric encoding location across trials (plus a random jitter of ±1 vu), which we refer to as “stable objects.” The other three objects have different (i.e., random) encoding locations across trials, and we refer to these objects as “unstable objects.” During each encoding, one stable and one unstable object is encountered. Objects are placed in the environment at least 1.5 vu away from the nearest landmark (tree). Stable objects are placed at least 10 vu away from other stable objects and at least 5 vu away from the center of the environment. Unstable objects are placed at least 10 vu away from other objects on the same trial and at least 1 vu away from all other objects on all trials. Following each fourth round, a score screen is displayed, providing participants with feedback on their points per trial and the number of trials completed.

### 2.3. Participants

We performed this study in two phases, resulting in two cohorts of participants. We used cohort 1 to explore various behavioral effects during the task, and we used cohort 2 to test whether these effects were similar in an independent group of participants. After analyzing cohort 1, we registered our findings on the Open Science Framework (https://osf.io/xs7g4). We then analyzed cohort 2. In the figures and Supplemental Information, we present the results of all analyses for the two cohorts separately, as well as for the two cohorts combined (“Cohorts 1 and 2”). In the main text, we describe only the results from the combined sample for brevity and clarity reasons.

Both cohorts comprised a sample of *n* = 32 healthy young participants. The mean age of the participants in cohort 1 was 25 ± 3 [mean ± standard deviation (SD)] with 17 females. The mean age of the participants in cohort 2 was 25 ± 4 (mean ± SD), with 17 females. Additionally, to optimize statistical power in the age and gender analyses, we included another group of *n* = 16 participants that we excluded from cohort 2 in the process of data collection because, in this group, the bike handles were accidentally not visible during encoding [age: 27 ± 8 (mean ± SD), 7 females].

Participants with neurological or psychiatric disorders were excluded from the study. Prior to participation, written informed consent was obtained, and participants were informed of their right to withdraw at any time. We recruited participants through flyers and verbal communication. Participants received 30 euros as compensation for their participation.

### 2.4. Software & Equipment

We created the Garden Game task using Unity (Unity Technologies, San Francisco, CA, USA; version 2021.3.14f1) under Windows 11 with the *C*# programming language. We provide the source-code version and the build version of the Garden Game for download online, along with detailed information on using the Garden Game (https://github.com/BonnSpatialMemoryLab/GardenGameTask). During the task, the Garden Game generates a log file containing various task and behavioral data, including participants’ spatial locations, orientations, and scores.

Together with the Garden Game, we provide a script in MATLAB (The MathWorks, Natick, MA, US) for creating custom trial-configuration files. These text files are read by the Garden Game during runtime and determine the structure of the task. Users can flexibly adjust this MATLAB script or change the trial-configuration files by hand to modify the Garden Game structure according to their needs.

We also provide two options for synchronizing the behavioral Garden Game data with other types of data such as neural data. The first synchronization option makes use of a MATLAB script that sends USB triggers to, for example, a neural recording system using functions from Psychtoolbox (Kleiner et al., 2007). In synchrony with these USB triggers, the MATLAB script simulates button presses that the Garden Game logs as triggers. The second synchronization option displays a small rectangle in the lower-right corner of the laptop screen. Changes in its color from black to white can be detected by a phototransistor, with its output connected to the analog channels of another recording device. These options will help users synchronize neural activity and other measures with the behavioral data from the Garden Game.

To perform eye tracking during the Garden Game, users may connect different types of eye trackers by Tobii (Tobii AB, Stockholm, Sweden) to the experiment laptop. Eye-tracking data is then included in the behavioral log file.

The experimental setup of this study comprised a laptop with a resolution of 2560 × 1600 pixels, operating at 60 frames per second for the first cohort and 165 frames per second for the second cohort. Participants navigated the virtual environment using a gamepad.

### 2.5. Eye tracking

In the second cohort, we performed eye tracking during the Garden Game, using the Tobii Pro Spark eye tracker (60 Hz temporal resolution). Participants with eye tracking received the same task instructions as those without eye tracking, and they were not given any additional instructions regarding their viewing behavior.

To investigate whether viewing behavior during encoding correlated with memory performance during retrieval, we quantified the time periods participants spent looking at specific environmental features (such as the boundaries and the corners) during encoding. To facilitate comparisons across trials, we normalized these time periods by the durations of the encoding periods containing valid eye tracking data. To determine the extent to which participants visually explored the environment during each encoding period, we estimated their “gaze coverage” of the environment. To this end, we divided the environment into 100 bins and labeled a bin as explored if the participant looked at it continuously for at least 250 ms. Gaze coverage was then defined as the proportion of bins explored by the participant. For example, gaze coverage was 50% if the participant looked at 50 of the 100 bins on a given trial continuously for at least 250 ms. To estimate the time participants focused on the relationship between their starting position and the object’s position, we quantified the time periods participants viewed the ground within an elliptic area between the object and the participant’s starting position (diameter of 1 vu).

### 2.6. Data exclusion

The initial training trial and all trials with one or more recall periods longer than 90 seconds were excluded from all subsequent analyses. This included trial 38 from participant 2 and trial 54 from participant 16.

### 2.7. Statistical analysis

All analyses were performed using Python (version 3.11.5). We mainly used linear mixed models (LMMs), with participants as random effects to account for individual variability. The dependent variable was allocentric or egocentric memory performance. All continuous predictors were mean-centered for LMMs with interaction terms to ensure interpretability (Lorah, 2020). If not otherwise specified, the LMMs included a single predictor of memory performance. For age and gender analyses, both predictors were included together in a single LMM. We computed the LMMs using the MixedLM.from formula function from the statsmodels library (Seabold and Perktold, 2010). All results of the LMMs analyzing allocentric and egocentric memory performance are presented in Tables S1 and S2. Statistical results not obtained through LMMs are reported in Table S3. To examine how allocentric and egocentric directions modulate memory performance, we used Friedman tests. This was followed by Wilcoxon signed-rank tests as post-hoc tests, with Sidak correction applied to correct for multiple comparisons (Tables S4, S5, and S6). In all analyses, we used the standard alpha-level of *α* = 0.05 to determine significant differences from those expected under the null hypothesis.

To examine learning curves related to individual objects, we fitted a Weibull curve to the trial-wise memory performance values using nonlinear least squares, separately for each object. The Weibull function is defined as

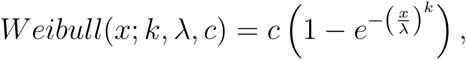

with *x* being the trial indices, *k* the abruptness of onset, *λ* the latency, and *c* the asymptote (Gallistel et al., 2004). The initial values assigned to the parameters were 2 for *k*, 10 for *λ*, and 1 for *c*. We constrained the three parameters to the possible ranges of [0.01, 0.01, 0] for the lower bounds and [100, 20, 1] for the upper bounds. In addition, the maximum number of function evaluations was set to 10,000 to allow sufficient time for the algorithm to converge. The optimized model parameters were then used to generate predicted values.

To identify the point at which learning occurred most strongly, which we refer to as the change point (Gallistel et al., 2004), we calculated a *t*-test for each trial *i* between the *y*-values of the Weibull function up to and including trial *i* to those after trial *i*. When the variance of the *y*-values was very small (smaller than 0.001), the fitted Weibull function was a completely or nearly horizontal line. This flatness indicates the absence of a learning process, and consequently, no change point was identified. After calculating this *t*-value for each trial *i*, we compared it against 1,000 surrogate *t*-values obtained by shuffling the raw performance scores, fitting the Weibull function, and repeating the same procedure as for the empirical data. We then selected the trial *i* with the highest proportion of surrogate *t*-values being smaller than the observed *t*-value (corresponding to the highest surrogate-corrected *t*-value) as the change point. A high value (close to 1) indicates that the observed change is significantly larger than would be expected by chance, while a low value (close to 0) indicates that the change is negligible.

Boxplots show the median across participants as thick black line, the interquartile range as box, minimum and maximum as whiskers, and outliers as dots. Outliers were identified as elements that exceeded the third quartile by more than 1.5 interquartile ranges or fell below the first quartile by more than 1.5 interquartile ranges. Regression plots were generated using the ggplot2 (Wickham, 2016) package in R, with model predictions and confidence intervals derived using effects (Fox and Weisberg, 2018, 2019) and regression models fitted with lme4 (Bates et al., 2015). The thick lines in the regression plots represent the predicted fit based on the LMM, and shaded areas indicate the 95% confidence intervals of these predictions.

## 3. Results

### 3.1. Task and basic behavioral results

We aimed at identifying various factors influencing allocentric and egocentric spatial memory using the Garden Game task (Figure 1; Materials & Methods). In brief, on each trial, participants navigated a virtual garden environment with a square boundary and encountered two out of six objects (animals) at different locations within the environment (two encoding periods per trial; Figure 1A). Participants were instructed to remember the positions of these objects with respect to the environment (allocentrically) and relative to their starting position (egocentrically). During retrieval, participants were asked to recall the locations of the two objects in abstract allocentric and egocentric reference frames (four individual recalls per trial; Figure 1, B and C). After each retrieval, participants immediately received spatial feedback regarding the accuracy of their response by viewing the correct location of the object, providing them with the opportunity of improving their response accuracy in subsequent retrievals.

Each participant completed 60 trials, with a total duration of 66.19 ± 1.56 minutes on average [mean ± standard error of the mean (SEM); Figure 2, A and B]. As we assigned feedback points using an adaptive procedure to enhance participants’ task engagement, they tended to receive about 28–32 out of 40 points per trial (Figure 2C). The random yet uniformly distributed starting positions, starting orientations, and object locations resulted in comprehensive environmental coverage of the navigation paths during encoding, as well as extensive coverage of the allocentric and egocentric reference frames during retrieval (Figure 2, D–G).

### 3.2. Associations of time and object stability with memory performance

Across the 60 trials, participants repeatedly encountered six different objects, with three of them appearing in approximately the same location across trials (referred to as “stable objects”) and three of them having different locations across trials (“unstable objects”; Figure 3A). This manipulation allowed us to investigate whether allocentric and egocentric memory performance varied as a function of “object stability” (i.e., whether the objects had a similar allocentric location during encoding across trials).

**Figure 3:**
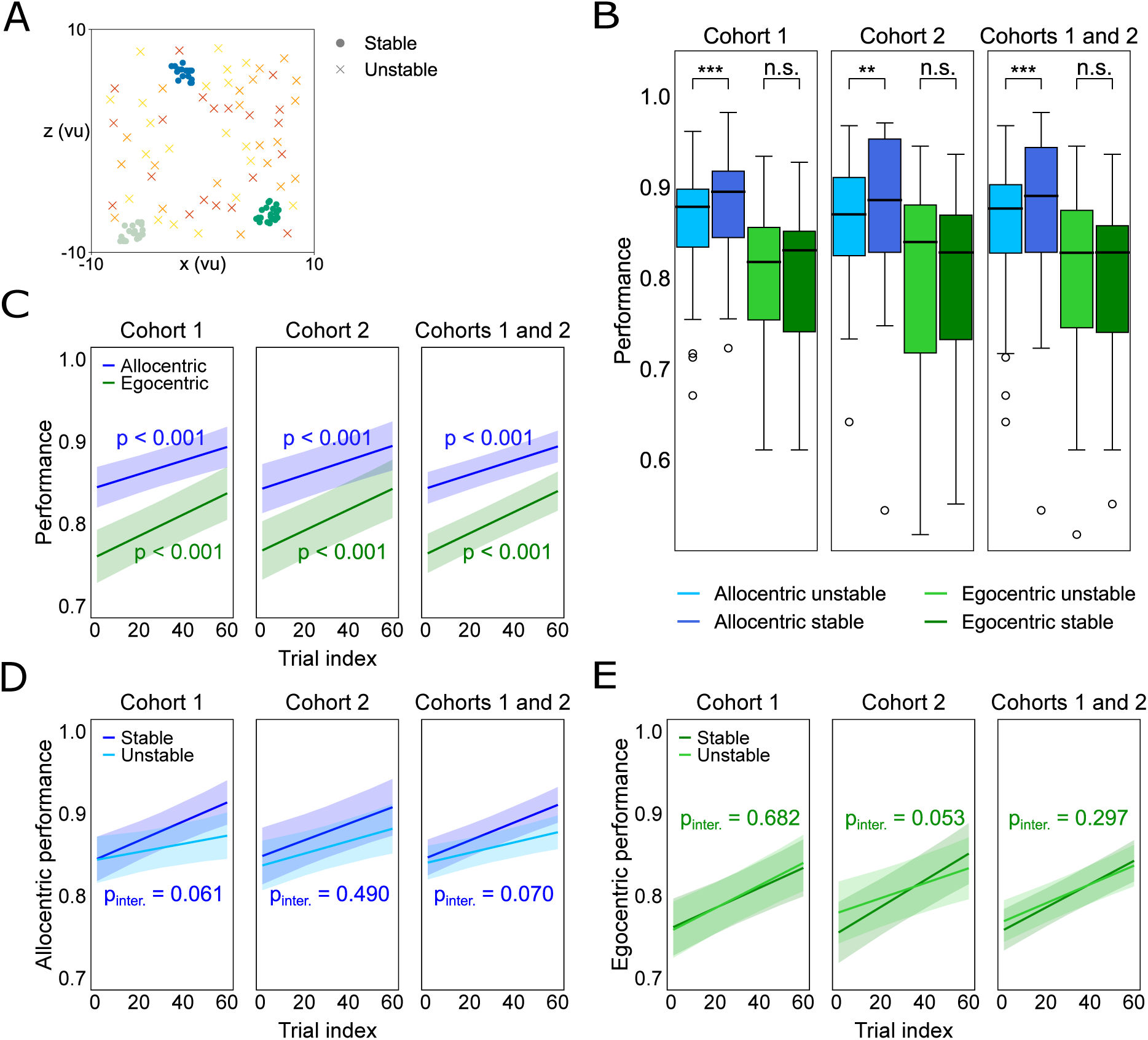
Associations of time and object stability with memory performance. (A) In each session, the Garden Game task features six different objects (animals), three of which are presented in the same location (plus a small jitter of ±1 vu; “stable objects”) and three of which are presented in different locations across all encoding periods (“unstable objects”). This design enables the investigation of one-shot learning (unstable objects) and of iterative learning (stable objects). The figure shows the locations of both stable (dots) and unstable (crosses) objects across all trials for an example participant. (B) Memory performance as a function of recall type (allocentric versus egocentric recall) and object stability (stable versus unstable objects). Allocentric memory performance was generally higher than egocentric memory performance, and allocentric memory performance was better for stable than for unstable objects. Boxplots show the median across participants as thick black line, the interquartile range as box, minimum and maximum as whiskers, and outliers as dots. (C) Allocentric and egocentric memory performance increased over time. (D) Allocentric memory performance increased over time. Performance tended to increase more strongly for stable than for unstable objects (see *p_inter_*). (E) Egocentric memory performance increased over time. Regression plots show predicted fits as thick lines, and shaded areas represent 95% confidence intervals. \*\**p* < 0.01; \*\*\**p* < 0.001; n.s., not significant; vu, virtual units.

In general, memory performance was higher for allocentric retrieval than for egocentric retrieval (paired *t*-test, *t* = 8.566, *p* < 0.001, *n* = 64), and participants with better allocentric memory performance also showed better egocentric memory performance (Pearson correlation, *r* = 0.734, *p* < 0.001, *n* = 64). Allocentric memory performance was better for stable objects than for unstable objects (LMM, *z* = 4.492, *p* < 0.001, *n* = 64), whereas egocentric memory performance was similar between stable and unstable objects (LMM, *z* = −0.505, *p* = 0.613, *n* = 64; Figure 3B). Accordingly, an LMM with object stability and type of memory as predictors showed a significant interaction, indicating that the effect of object stability was specific to allocentric memory (LMM, *z* = −3.419, *p* = 0.001, *n* = 64). These findings indicate that participants successfully used the consistent spatial relationships between the stable objects and the environment for allocentric but not egocentric retrieval.

Allocentric memory performance increased over time (LMM, *z* = 6.748, *p* < 0.001, *n* = 64; Figure 3C), both for stable and unstable objects (LMM, *z* = 6.074, *p* < 0.001, *n* = 64 and *z* = 3.498, *p* < 0.001, *n* = 64, respectively). When testing for a difference in allocentric memory improvement over time as a function of object stability, we observed a trend toward stronger improvement for stable as compared to unstable objects (LMM, *z* = 1.809, *p* = 0.070, *n* = 64; Figure 3D). We also observed a clear increase in memory performance for egocentric retrieval (LMM, *z* = 9.629, *p* < 0.001, *n* = 64; Figure 3C), with similar increases for stable objects as compared to unstable objects (LMM, *z* = 1.042, *p* = 0.297, *n* = 64; Figure 3E).

Together, these results demonstrate that time (experience) and object stability are major influencing factors of allocentric and egocentric memory recall. The effect of time is common to allocentric and egocentric memory, whereas the effect of object stability is specific to allocentric memory.

### 3.3. Object-specific learning

In the previous subsection, we performed learning analyses on the relationship between memory performance and time by collapsing across different objects. Next, we performed such learning analyses for each object individually, which we considered to reveal additional insights into the learning dynamics of allocentric and egocentric spatial memory (Gallistel et al., 2004).

By visually inspecting the object-specific learning curves, we observed approximately four different types of learning that we termed “abrupt learning,” “gradual learning,” “no learning at high performance,” and “no learning at low performance.” Abrupt learning was characterized by a rapid improvement in performance from one trial to the next (e.g., Figure 4A, first column). In gradual learning, performance improved steadily over time (Figure 4A, second column). Some learning curves were not indicative of learning. Their performance values were either near perfect from the beginning (no learning at high performance; Figure 4A, third column) or remained randomly distributed throughout the task (no learning at low performance; Figure 4A, fourth column).

**Figure 4:**
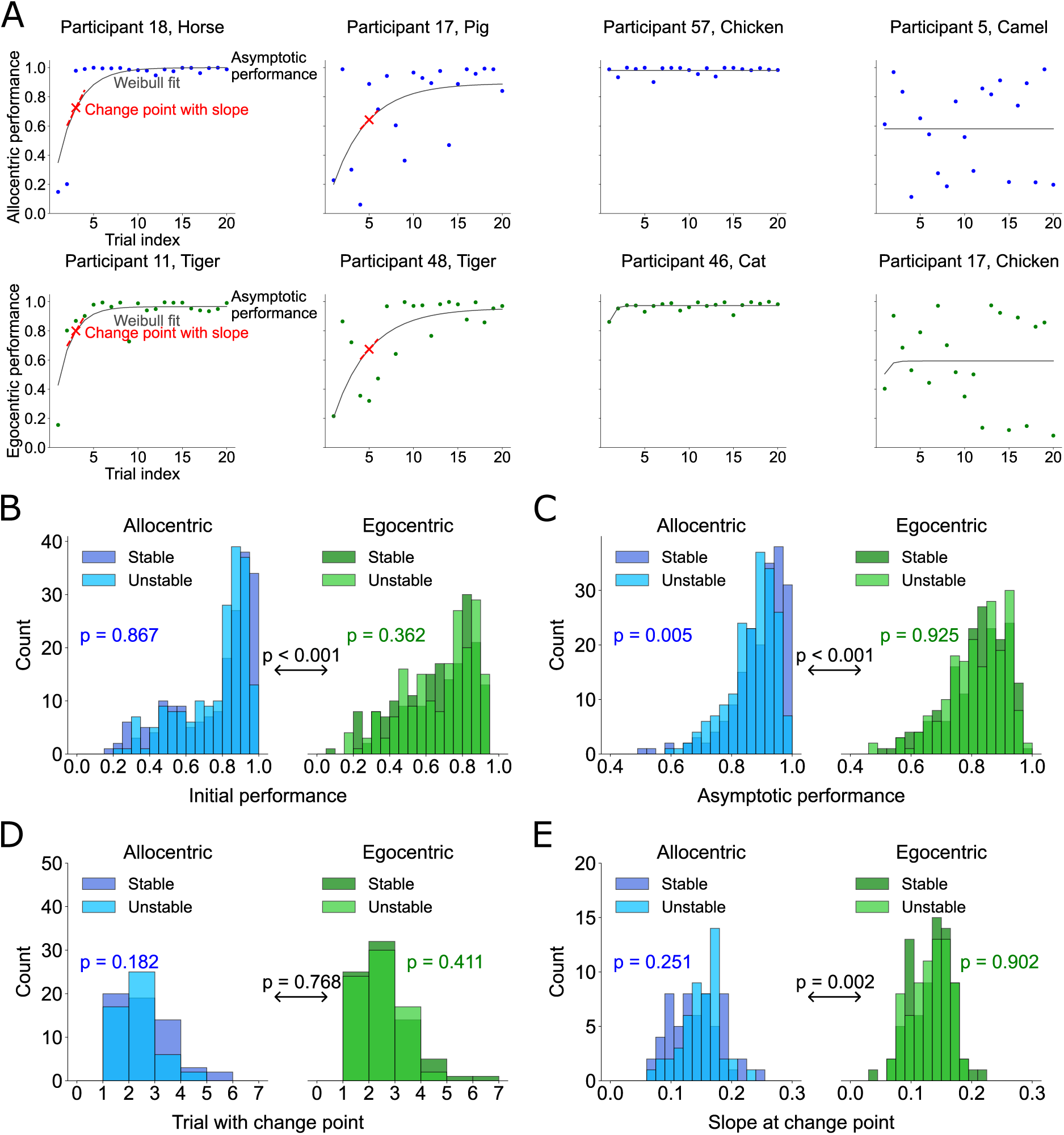
Object-specific learning. (A) Examples of object-specific learning curves. The first row corresponds to allocentric retrieval and the second row to egocentric retrieval. The first column represents abrupt learning and the second column gradual learning. The third column shows no learning with high performance from the beginning, and the fourth row shows no learning with poor performance throughout the task. The dots show the raw memory performance in each trial, while the gray line corresponds to the fitted Weibull function. The red cross indicates the change point, and the dashed line corresponds to the slope at the change point. Asymptotic performance is the *c* value from the fitted Weibull function. (B) Histograms of the initial performance values for allocentric and egocentric learning curves. Initial performance was higher for allocentric than for egocentric learning curves. (C) Histograms of the asymptotic performance values for allocentric and egocentric learning curves. Asymptotic performance was higher for allocentric than for egocentric learning curves. Regarding allocentric learning curves, asymptotic performance was higher for stable than for unstable objects. (D) Histograms of the trial indices of the change points for allocentric and egocentric learning curves. (E) Histograms of the slopes at the change points for allocentric and egocentric learning curves. Slopes were higher for allocentric than for egocentric learning curves.

To quantify each object-specific learning curve, we fit a Weibull function to the memory performance values across trials (Gallistel et al., 2004). We extracted four characteristics from the fitted learning curves: the initial performance, the asymptotic performance, the change point where learning occurred most strongly, and the slope of the learning curve at the change point (Materials & Methods). In brief, we defined initial performance as the fitted performance at the first trial, and we defined asymptotic performance as the parameter *c* of the fitted Weibull function. We identified the change point as the index of the trial leading to the strongest improvement in memory performance when comparing later versus earlier trials, and we estimated the slope as the change in performance at the change point by calculating the numerical gradient at the change point.

We found that initial and asymptotic performance was higher for allocentric than for egocentric learning curves (*t*-tests, *t* = 6.864, *p* < 0.001, *n* = 384 and *t* = 9.518, *p* < 0.001, *n* = 384, respectively; Figure 4, B and C), indicating that participants were better at allocentric as compared to egocentric memory recall throughout the task and early on. For allocentric learning curves, we found higher asymptotic performance values for stable than for unstable objects (*t*-test, *t* = 2.836, *p* = 0.005, *n* = 192; Figure 4C), with no difference in initial performance (*t*-test, *t* = −0.167, *p* = 0.867, *n* = 192; Figure 4B). For egocentric learning curves, participants showed similar initial and asymptotic performance between stable and unstable objects (*t*-tests, *t* = −0.913, *p* = 0.362, *n* = 192 and *t* = 0.095, *p* = 0.925, *n* = 192, respectively). These results align with and extend the above-mentioned effects of time and object stability on memory performance, showing that a consistent relationship between the object and the environment correlates with better allocentric memory performance and that this effect strengthens over time.

Across object-specific learning curves, we obtained more change points for egocentric learning curves than for allocentric learning curves (*χ*^2^-test, *χ*^2^ = 10.276, *p* = 0.001, *n* = 384), presumably because more allocentric learning curves fell into the category of no learning at high performance. We found similar numbers of change points between stable and unstable objects for both allocentric and egocentric memory (*χ*^2^-tests, allocentric: *χ*^2^ = 0.631, *p* = 0.427, *n* = 192; egocentric: *χ*^2^ = 0.175, *p* = 0.676, *n* = 192). Change-point trials occurred early, between object-specific trials 1 and 7, with no differences observed between allocentric and egocentric learning curves (*t*-test, *t* = −0.295, *p* = 0.768, *n_allo_* = 108, *n_ego_* = 151; Figure 4D) or between stable and unstable objects (*t*-tests, allocentric: *t* = 1.344, *p* = 0.182, *n_stable_* = 58, *n_unstable_*= 50; egocentric: *t* = 0.825, *p* = 0.411, *n_stable_* = 78, *n_unstable_*= 73; Figure 4D).

The slope at the identified change points was higher for allocentric than for egocentric learning curves (*t*-test, *t* = 3.128, *p* = 0.002, *n_allo_* = 108, *n_ego_*= 151; Figure 4E), indicating more rapid shifts in allocentric versus egocentric memory performance. There were no differences between stable and unstable objects (*t*-tests, allocentric: *t* = −1.156, *p* = 0.251, *n_stable_* = 58, *n_unstable_* = 50; egocentric: *t* = 0.123, *p* = 0.902, *n_stable_* = 78, *n_unstable_* = 73; Figure 4E).

Together, these results suggest that objects with a consistent relationship to the environment become easier to remember allocentrically over time, resulting in a significant difference in asymptotic performance for stable versus unstable objects. When learning occurred, it was more pronounced in allocentric memory, reflected by higher slopes at the change points of allocentric versus egocentric learning curves.

### 3.4. Associations of environmental features with memory performance

The navigable area of the virtual environment comprised a grassy area with an edge length of 20 vu, surrounded by a square boundary (fence). The north boundary was highlighted in black color as compared to the other three boundaries. Three trees located at the centers of three quarters of the environment served as intramaze landmarks (Figure 5A).

**Figure 5:**
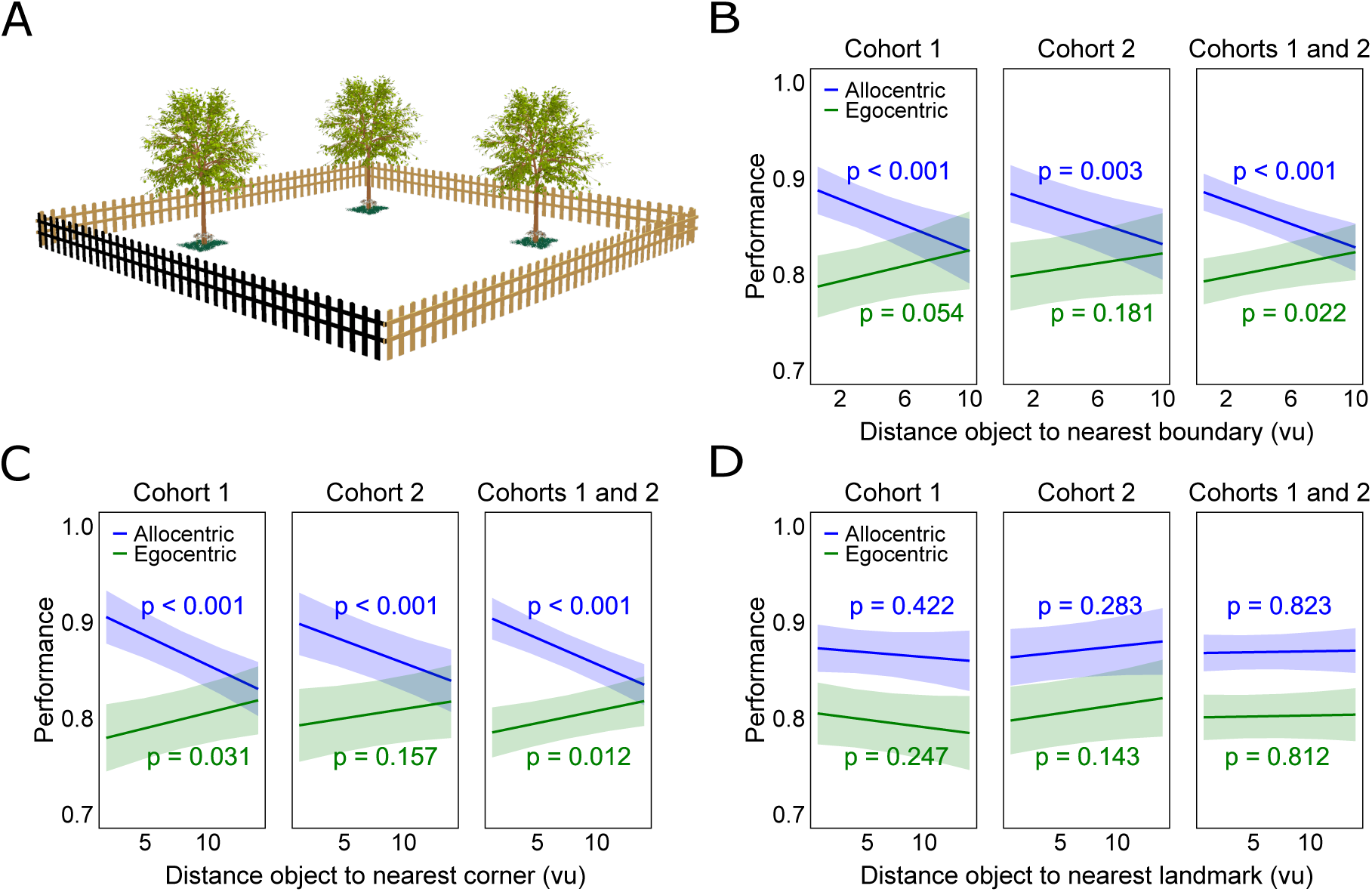
Associations of environmental features with memory performance. (A) The virtual environment of the Garden Game task includes four boundaries (fences). The north boundary is highlighted in black color, providing a strong orientation cue, and the other boundaries are shown in brown color. Three trees are placed within the environment as intramaze landmarks. The landmark configuration is randomized across participants. Illustration created using Autodesk Maya 3D 2024. (B) Relationship between the object’s distance toward the nearest boundary and memory performance. The closer the object to the nearest boundary, the better allocentric memory performance. The closer the object to the nearest boundary, the worse egocentric memory performance, which is a side effect of the object’s distance to the participant’s starting position (see Figure 6B). (C) Relationship between the object’s distance to the nearest corner and memory performance. The closer the object to the nearest corner, the better allocentric memory performance. The closer the object to the nearest corner, the worse egocentric memory performance, which is again a side effect of the object’s distance to the participant’s starting position (see Figure 6B). (D) No relationship between the object’s distance to the nearest intramaze landmark and memory performance. Regression plots show predicted fits as thick lines, and shaded areas represent 95% confidence intervals. vu, virtual units.

For allocentric retrieval, memory performance was significantly better for objects that were closer to the boundary (LMM, *z* = −4.592, *p* < 0.001, *n* = 64; Figure 5B). This effect was strongest for the north boundary (LMM, *z* = −5.166, *p* < 0.001, *n* = 64). Similarly, allocentric memory performance was better the closer the objects were to the corners (LMM, *z* = −5.618, *p* < 0.001, *n* = 64; Figure 5C).

Egocentric performance decreased for objects that were closer to the borders and corners (LMMs, *z* = 2.299, *p* = 0.022, *n* = 64 and *z* = 2.517, *p* = 0.012, *n* = 64, respectively; Figure 5, B and C), meaning that performance was better when the objects were nearer to the center of the environment. We reasoned that this result was a spurious side effect of the fact that objects closer to the participant’s starting position were associated with better egocentric memory performance (see subsection 3.5). Accordingly, this effect disappeared when we controlled for the distance between the object and the participant’s starting position on each trial (LMMs, *z* = 0.824, *p* = 0.410, *n* = 64 for borders and *z* = 0.531, *p* = 0.595, *n* = 64 for corners). Still, egocentric memory performance was better when the objects were closer to the north boundary (LMM, *z* = −3.940, *p* < 0.001, *n* = 64), even when controlling for the distance between the object and the participant’s starting position (LMM, *z* = −4.032, *p* < 0.001, *n* = 64).

Regarding the distance between the object and its nearest intramaze landmark (tree), we did not observe any association with allocentric or egocentric memory performance (LMMs, allocentric: *z* = 0.223, *p* = 0.823, *n* = 64; egocentric: *z* = 0.238, *p* = 0.812, *n* = 64; Figure 5D). We believe this null effect occurred because the intramaze landmarks were not visible during either allocentric or egocentric retrieval.

These results show that the boundaries and corners of the garden environment served as important determinants of allocentric retrieval. Additionally, the major visual orientation cue (the distinctively colored north boundary) supported both allocentric and egocentric memory retrieval.

### 3.5. Associations between starting positions and memory performance

At the beginning of each encoding period, participants were located at a specific allocentric starting position with a specific allocentric orientation, serving as the reference point for egocentric retrieval (Figure 6A). To-be-remembered objects were then presented at particular allocentric positions within the environment, thus appearing at specific egocentric distances and directions relative to the participants’ starting position. Hence, we aimed at understanding how egocentric distances and directions as well as allocentric starting orientations were associated with allocentric and egocentric performance (Figure 6).

**Figure 6:**
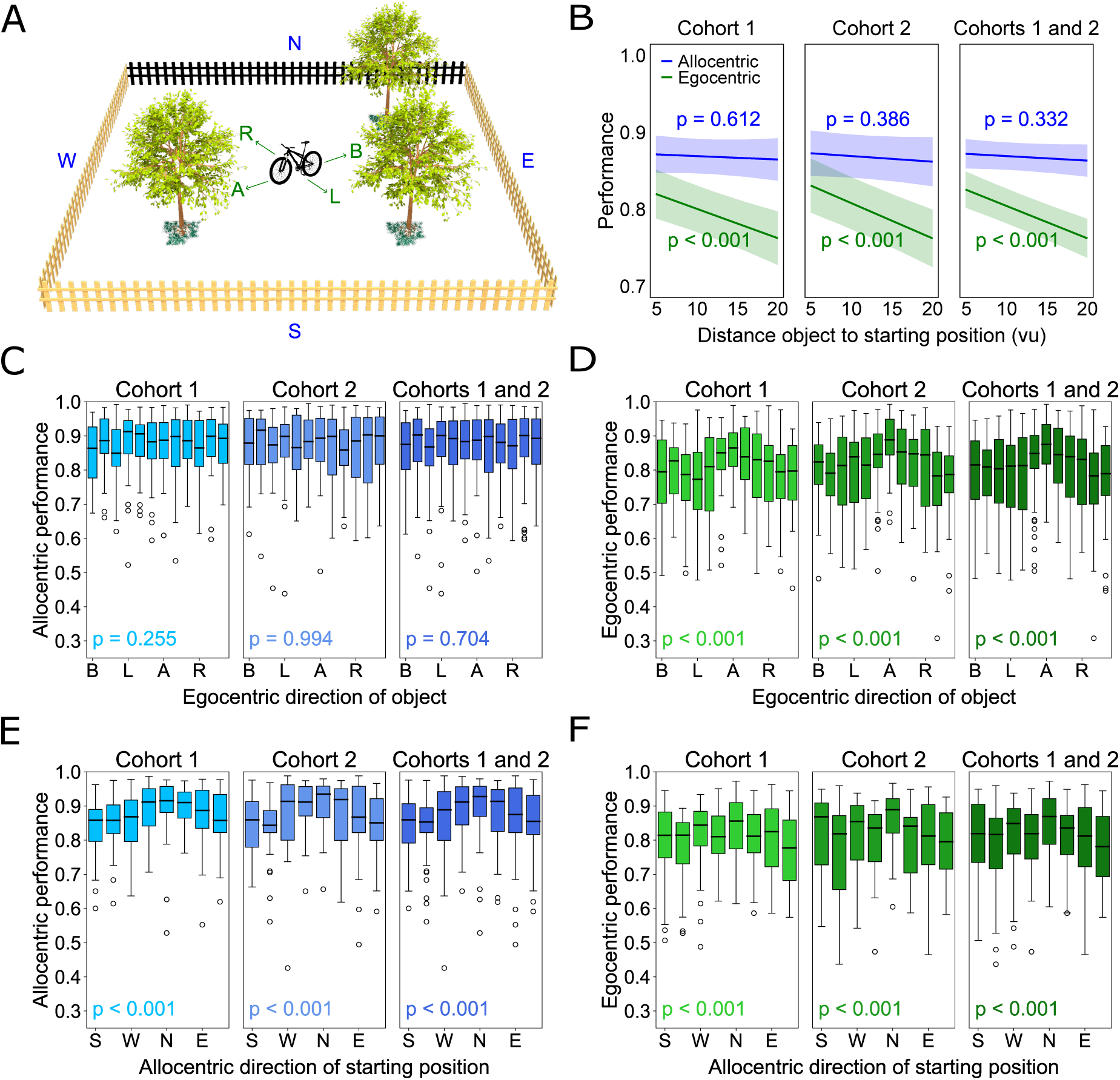
Associations between properties of the starting positions and memory performance. (A) Illustration of allocentric and egocentric reference frames at an example starting position, overlaid onto the virtual environment. Participants ride a bike and see the bike handles from their first-person perspective during encoding (Figure 1A). Illustration created using Autodesk Maya 3D 2024. (B) Relationship between the object’s distance to the starting position and memory performance. The smaller the distance, the better egocentric memory performance. (C) Relationship between allocentric memory performance and the egocentric direction between the object and the starting position. Performance did not vary as a function of egocentric direction. (D) Relationship between egocentric memory performance and the egocentric direction between the object and the starting position. Egocentric memory performance was higher when the object was ahead of the participant’s starting position. (E) Relationship between allocentric memory performance and the allocentric direction of the starting position. Performance was higher when the participant’s starting position was oriented toward north. (F) Relationship between egocentric memory performance and the allocentric direction of the starting position. Performance was higher when the participant’s starting position was oriented along the two cardinal axes of the environment. Boxplots show the median as thick black line, the interquartile range as box, minimum and maximum as whiskers, and outliers as dots. Regression plots show predicted fits as thick lines, and shaded areas represent 95% confidence intervals. A, ahead; B, behind; L, left; R, right. N, north; E, east; S, south; W, west. vu, virtual units.

Regarding egocentric distances, we found that egocentric memory performance was better for objects closer to the participant’s starting position (LMM, *z* = −6.615, *p* < 0.001, *n* = 64). This effect was not present for allocentric memory performance (LMM, *z* = −0.969, *p* = 0.332, *n* = 64; Figure 6B).

To examine the effects of egocentric directions, we divided the directions into twelve bins: ahead (A), left (L), right (R), behind (B), and two additional bins between each pair. We found significant differences in egocentric memory performance depending on the egocentric direction between the objects and participants’ starting orientation (Friedman test, *F* = 11.396, *p* < 0.001, *n* = 64). Egocentric performance was particularly good when the object was positioned ahead of the participant’s starting orientation (post-hoc comparisons of ahead with the other nine bins not facing ahead, all *W* ≤ 406, all *p_corr._* < 0.002, Sidak corrected for 66 comparisons; Figure 6D; Table S4). There were no differences in allocentric memory performance as a function of egocentric directions (Friedman test, *F* = 0.736, *p* = 0.704, *n* = 64; Figure 6C). Hence, egocentric memory performance—but not allocentric performance—was strongly modulated by the egocentric relationships of the objects relative to participants’ starting positions.

To analyze the effects of allocentric starting orientations on memory performance, we categorized them into eight bins: north (N), north-east (NE), east (E), south-east (SE), south (S), south-west (SW), west (W), and north-west (NW). This showed that allocentric memory performance varied significantly as a function of the allocentric starting orientation (Friedman test, *F* = 9.255, *p* < 0.001, *n* = 64). Specifically, performance during allocentric retrieval was significantly higher when the participant’s starting orientation faced a northward direction compared to any southward direction (all *p_corr._* < 0.012, with *p_corr._* = 0.058 for the comparison between NE and SE, Sidak corrected for 28 comparisons; Figure 6E; Table S5). Visual inspection of the performance histograms showed a unimodal distribution with a preference for the northward orientation.

We also observed a significant modulation of egocentric memory performance by allocentric starting orientation (Friedman test, *F* = 11.360, *p* < 0.001, *n* = 64). Performance was increased when the participants’ starting position was oriented towards north compared to all other directions (all *p_corr._* < 0.040, Sidak corrected for 28 comparisons; Figure 6F; Table S6). In this case, however, visual inspection of the performance histograms suggested a quadrimodal distribution of performance values as a function of allocentric starting orientation, with preferences for the two cardinal axes of the environment (i.e., along the north–south and the east–west axes). Accordingly, egocentric memory performance was significantly higher when participants’ starting orientation was aligned with the cardinal axes as compared to when misaligned with them (LME for N/E/S/W versus NE/SE/SW/NW, *z* = 6.075, *p* < 0.001, *n* = 64), in line with previous behavioral studies (Mou and McNamara, 2002; McNamara et al., 2003).

In summary, these findings support the idea that egocentric memory performance depends considerably on egocentric properties during encoding (i.e., the distances and directions of the objects relative to the initial starting positions). Its modulation by the alignment of participants’ allocentric orientation with the cardinal axes may be due to improved estimation of angular rotations when initially facing a boundary, or because of reduced mental rotation efforts when the egocentric cognitive map aligns with the environment’s privileged axes (McNamara et al., 2008; Gagnon et al., 2014). In contrast, allocentric memory performance was unaffected by egocentric properties during encoding. Instead, it was primarily determined by participants’ allocentric starting orientation, with a strong influence from the major visual cue (i.e., the distinctively colored north boundary).

### 3.6. Associations between spatial feedback and memory performance

During the retrieval phase, after participants responded, they immediately received spatial feedback indicating the correct position of the object in the abstract allocentric and egocentric reference frames (Figure 1, B and C). To determine whether this feedback facilitated subsequent recall, we compared memory performance values during retrieval periods with versus without feedback (i.e., after versus before feedback, respectively; Figure 7). We considered that positive effects of allocentric feedback on egocentric performance, and of egocentric feedback on allocentric performance, would indicate direct links between allocentric and egocentric cognitive maps. In contrast, no or negative feedback effects would be in line with a separation of allocentric and egocentric systems. Per trial, there were four individual recall periods, including two allocentric retrievals and two egocentric retrievals. The order of allocentric and egocentric retrieval alternated between trials, so that half of the trials started with allocentric retrieval and the other half started with egocentric retrieval. Retrieving the firstly or secondly encoded object first was randomized across all retrievals, making the retrieval results independent of the order in which participants encountered the objects during encoding (Figure 7A). By comparing particular retrieval positions with each other, this task design enabled us to investigate the effects of feedback on memory performance within allocentric/egocentric domains (i.e., allocentric feedback on allocentric performance, and egocentric feedback on egocentric performance) and across domains (allocentric feedback on egocentric performance, and egocentric feedback on allocentric performance). For example, comparing the first allocentric retrieval (at retrieval positions 1 or 3) with the second allocentric retrieval (at retrieval positions 2 or 4) allowed us to estimate the effects of allocentric feedback on allocentric performance.

**Figure 7:**
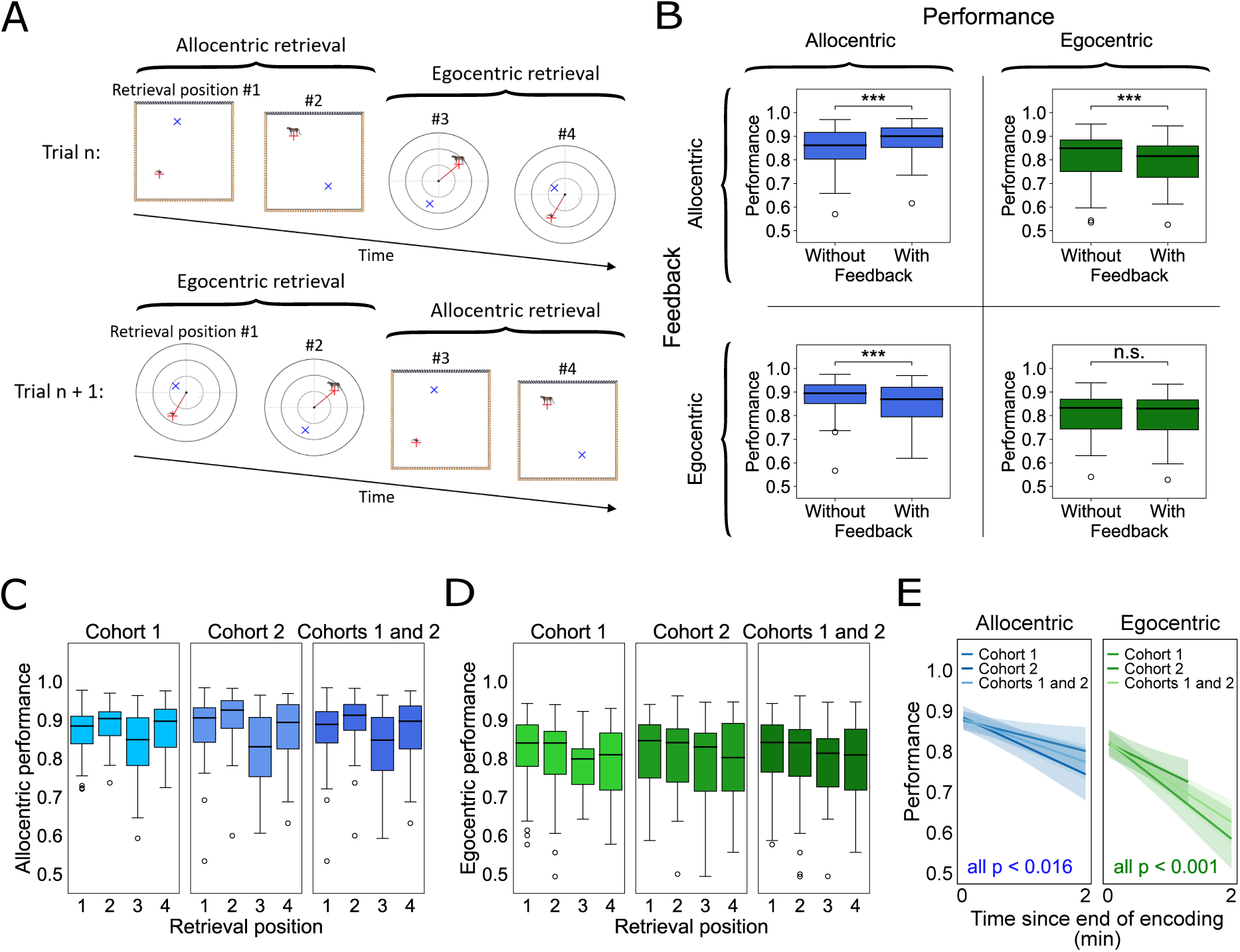
Associations between feedback and memory performance. (A) During each retrieval period, participants performed allocentric and egocentric retrieval for both objects from the preceding encoding period. After each response, participants immediately received feedback via a blue cross indicating the object’s correct location. The figure illustrates two retrieval periods, with allocentric (top) and egocentric retrieval first (bottom). The order of allocentric and egocentric retrieval alternated between trials. Plots show the time of feedback. Red cross, response position; blue cross, correct position of the object. (B) Effects of allocentric/egocentric feedback on allocentric/egocentric memory performance. Top left: Allocentric memory performance was better for allocentric retrieval after allocentric feedback (retrieval positions 2 and 4 versus 1 and 3), suggesting a positive effect of allocentric feedback on allocentric memory performance. Top right: Egocentric memory performance was worse for egocentric retrieval after allocentric feedback (retrieval positions 3 and 4 versus 1 and 2), suggesting no benefit of allocentric feedback on egocentric memory performance. Bottom left: Allocentric memory performance was worse after egocentric feedback (retrieval positions 3 and 4 versus 1 and 2), suggesting no benefit of egocentric feedback on allocentric memory performance. Bottom right: Egocentric memory performance was similar before and after egocentric feedback (retrieval positions 2 and 4 versus 1 and 3). Together with C–E, these findings suggest a major detrimental effect of time between encoding and retrieval on memory performance that can be fully compensated during allocentric retrieval by allocentric feedback and that can be partially compensated during egocentric retrieval by egocentric feedback. (C) Allocentric memory performance as a function of retrieval position, showing reduced performance for later retrieval (retrieval positions 3 and 4 versus 1 and 2) and better performance after allocentric feedback (retrieval positions 2 and 4 versus 1 and 3). (D) Egocentric memory performance as a function of retrieval position, showing reduced performance for later retrieval (retrieval positions 3 and 4 versus 1 and 2). (E) Association between memory performance and time between encoding and retrieval. Memory performance decreased for both allocentric and egocentric memory performance as more time elapsed since the end of encoding. Boxplots show the median across participants as thick black line, the interquartile range as box, minimum and maximum as whiskers, and outliers as dots. Regression plots show predicted fits as thick lines, and shaded areas represent 95% confidence intervals. \*\*\**p* < 0.001; n.s., not significant.

When testing the effects of feedback on performance within domains, we found that allocentric feedback had a positive effect on allocentric performance, as allocentric performance at retrieval positions 2 and 4 was better than allocentric performance at retrieval positions 1 and 3 (LMM, *z* = 7.837, *p* < 0.001, *n* = 64; Figure 7B). This positive effect survived the general trend of decreasing allocentric and egocentric performance values at successively later retrieval positions (Figure 7, C–E). Regarding egocentric retrieval, we found that egocentric feedback had no apparent effect on egocentric performance as egocentric performance at retrieval positions 2 and 4 was similar to egocentric performance at retrieval positions 1 and 3 (LMM, *z* = −1.783, *p* = 0.075, *n* = 64; Figure 7B).

We interpret this as a stabilizing effect of egocentric feedback on egocentric performance as it reduced the negative impact of later retrieval positions on egocentric performance (Figure 7, D and E, right).

When investigating cross-domain effects, we found that allocentric feedback appeared to have a negative impact on egocentric performance as egocentric performance at retrieval positions 3 and 4 was worse than egocentric performance at retrieval positions 1 and 2 (LMM, *z* = −3.997, *p* < 0.001, *n* = 64). Similarly, egocentric feedback was associated with worse allocentric performance as allocentric performance at retrieval positions 3 and 4 was worse than at retrieval positions 1 and 2 (LMM, *z* = −6.498, *p* < 0.001, *n* = 64; Figure 7B). Hence, across allocentric and egocentric domains, spatial feedback did not reverse the negative effects of later retrieval positions on performance. These results are in line with a separation of allocentric and egocentric systems.

We performed additional analyses to further understand this pattern of results. To this end, we analyzed memory performance as a function of retrieval position, separately for allocentric and egocentric recall. Comparing each retrieval position to position 1, we found that allocentric memory performance was better at retrieval position 2 (LMM, *z* = 3.810, *p* < 0.001, *n* = 64), worse at retrieval position 3 (LMM, *z* = −6.363, *p* < 0.001, *n* = 64), and similar at retrieval position 4 compared to position 1 (LMM, *z* = 0.944, *p* = 0.345, *n* = 64; Figure 7C). In contrast, egocentric memory performance consistently decreased across retrieval positions 2, 3, and 4 when compared to retrieval position 1 (LMM, position 2: *z* = −1.969, *p* = 0.049, *n* = 64; position 3: *z* = −3.533, *p* < 0.001, *n* = 64; position 4: *z* = −4.089, *p* < 0.001, *n* = 64; Figure 7D).

These results help visualize the above-mentioned effects of feedback on performance, showing most clearly that performance generally decreased at later retrieval positions and that allocentric feedback at retrieval positions 1 and 3 improved allocentric performance at retrieval positions 2 and 4.

We reasoned that poorer performance at later retrieval positions stemmed from longer time periods between encoding and retrieval, allowing for forgetting. Accordingly, both allocentric and egocentric memory performance decreased significantly as a function of time since the end of the encoding period (LMM, allocentric: *z* = −4.659, *p* < 0.001, *n* = 64; egocentric: *z* = −7.107, *p* < 0.001, *n* = 64; Figure 7E). We obtained similar results when using the object-specific end times of encoding, showing a decline in egocentric and allocentric performance over time for both the firstly encoded object (LMM, allocentric: *z* = −2.721, *p* = 0.007, *n* = 64; egocentric: *z* = −3.935, *p* < 0.001, *n* = 64) and the secondly encoded object (LMM, allocentric: *z* = −3.692, *p* < 0.001, *n* = 64; egocentric: *z* = −6.541, *p* < 0.001, *n* = 64).

Overall, these results show a strong detrimental effect of time since encoding on recall performance. Allocentric feedback fully compensated for this time effect regarding allocentric recall (better allocentric performance for retrieval positions 2 and 4 as compared to 1 and 3), and egocentric feedback partially compensated for it with regard to egocentric recall (similar egocentric performance for retrieval positions 2 and 4 as compared to 1 and 3; Figure 7B). Across allocentric and egocentric domains, spatial feedback did not support performance.

### 3.7. Associations between viewing behavior and memory performance

We sought to examine whether participants’ viewing behavior during encoding correlated with their memory performance during retrieval. Hence, in cohort 2, we recorded eye tracking data at 60 Hz while participants performed the task (Figure 8; for cohort 1, eye tracking was not available). For each encoding period, we quantified how much time participants spent viewing particular aspects of the environment and tested for associations with memory performance during allocentric and egocentric retrieval.

**Figure 8:**
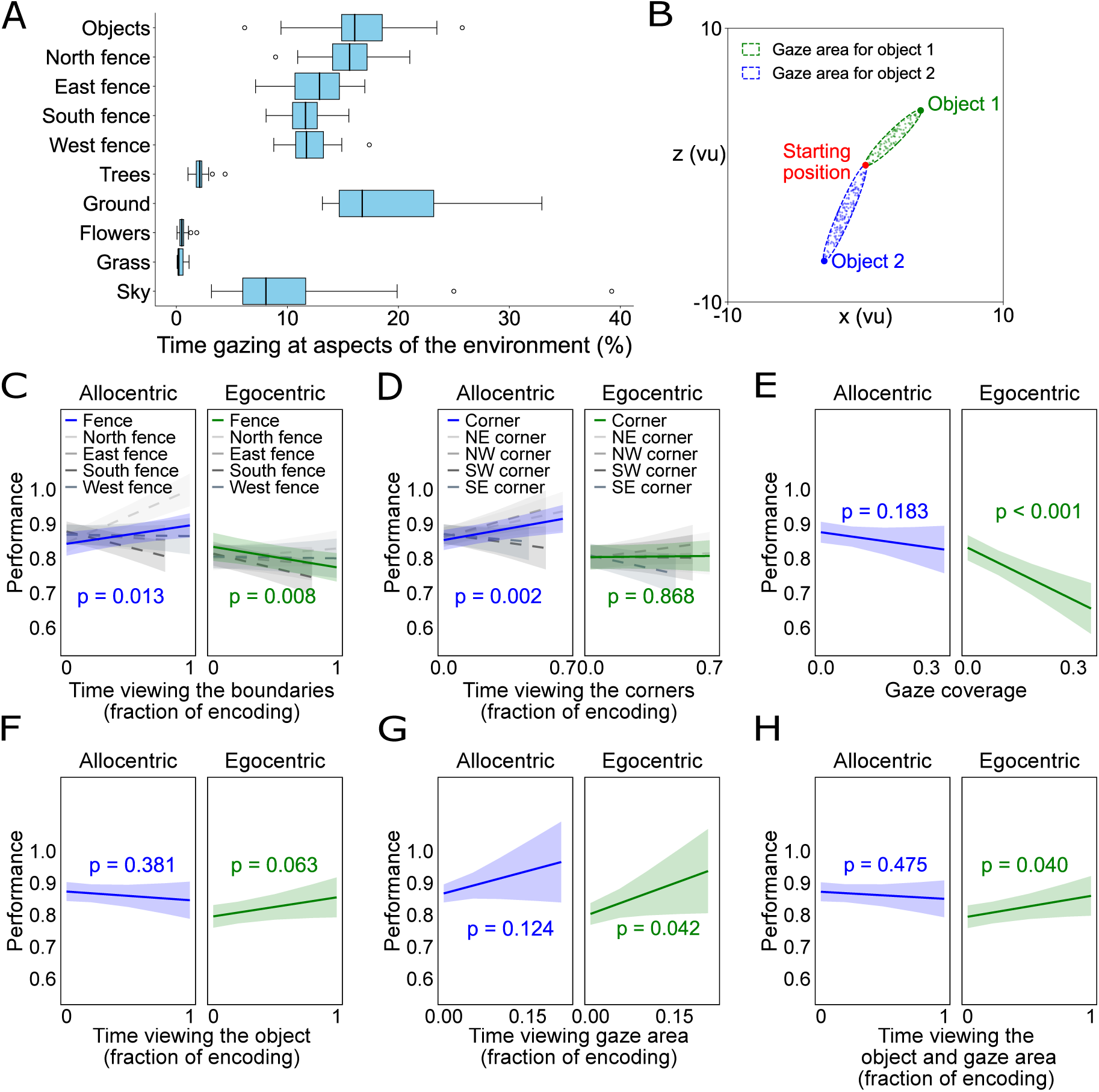
Associations between viewing behavior and memory performance (cohort 2 only). (A) Percentage of time gazing at different aspects of the virtual environment. Boxplots show the median across participants as thick black line, the interquartile range as box, minimum and maximum as whiskers, and outliers as dots. (B) Definition of the “gaze area.” For both objects during encoding, we estimated the time period participants spent gazing at an elliptic area on the ground between the object and the participant’s starting position (diameter of 1 vu). (C) Association between memory performance and percentage of time viewing the boundaries (fences). The longer participants gazed at the boundaries, the better allocentric memory performance and the worse egocentric memory performance. (D) Association between memory performance and percentage of time viewing the corners. The longer participants gazed at the corners, the better allocentric memory performance. (E) Association between memory performance and gaze coverage of the environment. The larger the area of the environment participants viewed during encoding, the worse egocentric memory performance. (F) Association between memory performance and percentage of time viewing the object. Egocentric memory performance tended to increase the longer participants gazed at the object. (G) Association between memory performance and percentage of time viewing the object-specific gaze areas. The longer participants viewed the gaze areas, the better egocentric memory performance. (H) Association between memory performance and percentage of time viewing the objects and their gaze areas. The longer the viewing time, the better egocentric memory performance. Regression plots show predicted fits as thick lines, and shaded areas represent 95% confidence intervals. vu, virtual units.

We first investigated the effects of viewing behavior that focused on the virtual environment, including the boundaries, the corners, and the ground. We hypothesized that focusing the gaze more on these environmental features would be correlated with better allocentric memory. Indeed, we found that trials in which participants spent more time viewing the boundaries during encoding were associated with better memory performance during allocentric retrieval (LMM, *z* = 2.473, *p* = 0.013, *n* = 32; Figure 8C). This effect was driven by a strong positive relationship between viewing the north boundary and allocentric memory performance (LMM, *z* = 6.663, *p* < 0.001, *n* = 32). In contrast, spending more time viewing the boundaries during encoding was generally associated with poorer egocentric memory performance during retrieval (LMM, *z* = −2.657, *p* = 0.008, *n* = 32), potentially because participants prioritized allocentric encoding of the object locations at the expense of egocentric memory in these trials.

We then examined the relationship between memory performance during retrieval and the time participants spent viewing the corners during encoding (defined as all 3D positions within ≤ 2 vu from a corner). Similar to our results on viewing the boundaries, longer viewing times of the environment corners were associated with better allocentric memory performance (*z* = 3.084, *p* = 0.002, *n* = 32). Egocentric memory performance showed no relationship with viewing the corners during encoding (*z* = 0.167, *p* = 0.868, *n* = 32; Figure 8D).

When we examined the impact of viewing large portions of the environment’s ground during encoding, quantified as “gaze coverage” (i.e., how many parts of the environment participants viewed continuously for ≥ 250 ms), we found no association with allocentric memory performance (LMM, *z* = −1.330, *p* = 0.183, *n* = 32; Figure 8E). This was unexpected, as we had anticipated greater gaze coverage to enhance allocentric memory by helping participants better situate the objects within the spatial scene. Furthermore, viewing larger portions of the ground during encoding was correlated with poorer egocentric memory performance (LMM, *z* = −4.523, *p* < 0.001, *n* = 32). This negative association may have arisen from higher gaze coverage being linked to participants making more turns, traveling longer distances, and taking more time during encoding—all of which tend to decrease egocentric memory performance.

Next, we investigated the effects of viewing behavior that focused on the participants’ relationship to the object, which we hypothesized to show positive associations with egocentric memory performance. We observed a trend for the effect that participants showed better egocentric memory performance during retrieval when they had spent more time viewing the objects during encoding (LMM, *z* = 1.859, *p* = 0.063, *n* = 32), with no effect on allocentric memory performance (LMM, *z* = −0.875, *p* = 0.381, *n* = 32; Figure 8F). Similarly, trials in which participants spent more time viewing the “gaze area” (i.e., an elliptic area with a diameter of 1 vu; Figure 8B) between the object and the participant’s starting position (which served as the reference point during egocentric retrieval) correlated with better egocentric memory performance (LMM, *z* = 2.034, *p* = 0.042, *n* = 32; Figure 8G), while there was no clear association with allocentric memory performance (LMM, *z* = 1.539, *p* = 0.124, *n* = 32). When combining both measures (i.e., viewing the object and viewing the gaze area), we again found a positive relationship to egocentric memory performance (LMM, *z* = 2.053, *p* = 0.040, *n* = 32; Figure 8H).

Taken together, these results show that gazing at important aspects of the environment including its borders and corners during encoding was linked to better allocentric memory performance during retrieval. Conversely, focusing the gaze on the to-be-remembered objects and their relationship to one’s own position correlated with better egocentric memory performance (though we note that these effects were relatively weak; Figure 8, F–H).

### 3.8. Effects of age and gender on memory performance

Age and gender are important determinants of spatial memory capabilities (Lester et al., 2017; Coutrot et al., 2018). We thus asked whether they were associated with allocentric and egocentric memory performance in the Garden Game. We performed these analyses separately for cohorts 1 and 2, and for both cohorts combined. Additionally, we included 16 participants initially excluded during cohort 2 data collection due to accidentally performing a slightly different version of the Garden Game (see Materials & Methods for details). Since our *a priori* design required using identical task versions across cohorts, these participants were excluded from the main analyses but we included them in the exploratory across-subjects age and gender analyses to increase statistical power. This “complete” sample thus comprised 80 participants, with an age range of 25 ± 5 years (mean ± SD), and 41 females.

We found that older age was generally related to poorer memory performance (Table 1), a finding that is consistent with previous research on age-related effects in spatial cognition (Kirasic, 1991; Park, 2000; Lester et al., 2017). We observed this age-related decline in memory performance for both allocentric and egocentric memory performance (LMMs, *z* = −2.241, *p* = 0.025, *n* = 80 and *z* = −2.126, *p* = 0.034, *n* = 80, respectively). We note though that this effect was modest, potentially due to the fact that the age ranges in our cohorts were relatively narrow (Table S3).

**Table 1:** Associations of age and gender with memory performance.

| Cohort | Allocentric Retrieval |  |  |  | Egocentric Retrieval |  |  |  |
| --- | --- | --- | --- | --- | --- | --- | --- | --- |
|  | Age |  | Gender |  | Age |  | Gender |  |
| | $z$ | $p$ | $z$ | $p$ | $z$ | $p$ | $z$ | $p$ |
| **Cohort 1** | 1.174 | 0.240 | -1.703 | 0.089 | -0.449 | 0.653 | 0.737 | 0.461 |
| **Cohort 2** | -2.321 | 0.020* | 2.325 | 0.020* | -1.650 | 0.099 | 2.528 | 0.011* |
| **Cohorts 1 and 2** | -1.570 | 0.116 | 0.932 | 0.351 | -1.743 | 0.081 | 2.445 | 0.014* |
| **Complete** | -2.241 | 0.025* | 1.067 | 0.286 | -2.126 | 0.034* | 2.411 | 0.016* |

The table presents the $z$ -scores and $p$ -values of an LMM with age and gender as predictors of allocentric and egocentric memory performance. Cohorts 1 and 2 comprise $n = 32$ participants each; the “complete” sample additionally includes $n = 16$ participants who completed a slightly different version of the Garden Game. \* $p < 0.05$ .

We found that male participants performed better during egocentric memory recall as compared to female participants (LMM, *z* = 2.411, *p* = 0.016, *n* = 80), similar to recent findings on gender differences in egocentric memory tasks (Fernandez-Baizan et al., 2019; Miola et al., 2023). Female and male participants performed similarly during allocentric recall (LMM, *z* = 1.067, *p* = 0.286, *n* = 80), though males tended to show better performance numerically. These results suggest that memory performance in the Garden Game is, to some extent, sensitive to gender differences, in line with previous studies on human spatial memory (Coluccia and Louse, 2004; Castelli et al., 2008; Boone et al., 2018; Munion et al., 2019; Clint et al., 2012; Coutrot et al., 2018).

## 4. Discussion

Recalling the locations of objects in the environment is an important aspect of everyday life (Postma et al., 2008). To this end, humans appear to use cognitive maps that provide mental representations of the environment (O’Keefe and Nadel, 1978; Ekstrom et al., 2018; Behrens et al., 2018; Bellmund et al., 2018). These cognitive maps can be allocentric, representing the objects’ locations relative to the external environment, or egocentric, representing them relative to the individual’s position and orientation (Klatzky, 1998; Burgess, 2006; Yeap, 2014; Ekstrom and Isham, 2017; Kunz et al., 2021; Kozhevnikov and Puri, 2023). Here, we introduced a new task, the Garden Game, to study the determinants of spatial memory when using allocentric and egocentric cognitive maps during memory recall (Figures 1 and 2). In each trial, participants navigated through a virtual garden environment during encoding and encountered two objects. Participants then recalled the object locations in abstract allocentric and egocentric reference frames relative to the environment and the initial starting position, respectively. We introduced these explicitly defined allocentric and egocentric reference frames to behaviorally probe participants’ use of allocentric and egocentric cognitive maps as directly as possible.

Our main results show that features of the relationship between objects and the environment during encoding (i.e., the objects’ distances to borders and corners) were correlated with memory performance during allocentric recall (Figure 5). In contrast, determinants of the relationship between the objects and participants’ starting positions (i.e., egocentric distances and directions) were linked to memory performance during egocentric recall (Figure 6). Furthermore, while there was a significant effect of allocentric starting orientations on both allocentric and egocentric recall, allocentric performance was unimodally driven by the major orientation cue (i.e., the distinctively colored north boundary) whereas egocentric memory varied as a function of participants’ alignment with the cardinal axes of the environment (Figure 6). Importantly, we found that feedback during allocentric retrieval improved allocentric performance, and that feedback during egocentric retrieval stabilized egocentric performance, but we did not observe positive effects of allocentric feedback on egocentric performance, and *vice versa* (Figure 7).

The observed distinct determinants of allocentric and egocentric memory performance, along with the lack of facilitation of memory performance through cross-domain feedback, indicate that allocentric and egocentric cognitive maps are, at least to some extent, separate from each other. We suggest that this interpretation is in line with the notion that allocentric and egocentric cognitive maps rely on distinct neural substrates comprising both allocentrically and egocentrically tuned neurons in various brain regions (Bicanski and Burgess, 2020; Wang et al., 2020; Alexander et al., 2023). The Garden Game offers a novel opportunity for studying and dissecting these systems at the behavioral and neural level, helping improve our understanding of human cognitive maps.

This study expands our knowledge of how characteristics of the environment, objects, and participants are correlated with allocentric and egocentric spatial memory performance. Relationships between the environment and the to-be-remembered objects were important for allocentric spatial memory. Participants showed better performance when the objects (animals) were closer to the boundaries and corners, supporting the idea that boundaries and corners are important for allocentric spatial memory (Hartley et al., 2004; Lee, 2017; Geva-Sagiv et al., 2023). At the neural level, this effect may be linked to the property of allocentric boundary-vector cells to represent shorter distances to boundaries more frequently than longer distances (Lever et al., 2009; Muessig et al., 2024). Eye tracking extended these findings by demonstrating that longer viewing times of the boundaries and corners were associated with better allocentric memory. In contrast, relationships between participants’ starting positions and the to-be-remembered objects were important for egocentric spatial memory. Specifically, shorter distances between the starting positions and the objects, as well as objects being positioned ahead of the starting positions, were associated with better egocentric memory. This observation may be connected to the neural finding that some types of egocentrically tuned cells prefer ahead directions and shorter distances (Wang et al., 2018; LaChance et al., 2019; Alexander et al., 2020; Kunz et al., 2021). Eye tracking extended these observations by showing improved egocentric memory performance when participants spent more time viewing the objects and their areas toward participants’ starting positions during memory encoding.

Investigating the effects of participants’ allocentric starting orientation provided additional insights into the distinct determinants of allocentric and egocentric spatial memory. Specifically, participants showed better allocentric memory when their starting position was oriented north, facing the boundary with a color distinct from the other three boundaries (i.e., the north boundary). Across all allocentric orientations, we observed a unimodal modulation of allocentric memory performance, with better performance when the starting position was oriented closer to north. With regard to egocentric memory, participants also showed better performance when participants started facing north. In this case, however, the data suggested a quadrimodal modulation, with better egocentric memory performance when participants’ allocentric starting orientation during encoding was aligned with any of the cardinal axes of the environment (i.e., along north–south or east–west). This finding may relate to the previous observation of better egocentric memory performance when remembering object locations from heading directions aligned with the intrinsic axes of the environment (Mou and McNamara, 2002; McNamara et al., 2003, 2008; Gagnon et al., 2014). In contrast to the modulations of both allocentric and egocentric memory performance by allocentric starting orientations, egocentric starting directions (e.g., whether an object was ahead of the participant) were associated with egocentric, but not allocentric, memory performance. Together, these results may imply some crosstalk between allocentric and egocentric computations with regard to allocentric starting orientations but not regarding egocentric starting orientations.

An important property of the objects was their stability over time, i.e., whether their allocentrically defined locations were similar across trials. This property had a major effect on allocentric memory performance, with stable objects (i.e., objects with similar allocentric encoding locations across trials) being associated with better performance than unstable objects (i.e., objects with different encoding locations across trials; Figure 3). In contrast, performance did not differ between stable and unstable objects during egocentric memory recall. We note though that our task did not have a condition in which object stability was defined relative to the egocentric frame of reference (i.e., objects having the same egocentric direction and distance toward the starting position across trials). Such a condition may be added in future studies, potentially demonstrating better egocentric performance for egocentrically-defined stable objects. Furthermore, participants tended to show stronger improvements in allocentric memory performance for stable versus unstable objects (Figure 3), resulting in significantly higher asymptotic allocentric memory performance for stable versus unstable objects (Figure 4). These results demonstrate that temporally stable relationships between an object and its surroundings facilitate allocentric but not egocentric memory, again suggesting that recall of egocentric information is not simply a readout-and-transformation of allocentric information, and *vice versa*.

In general, participants demonstrated strong learning for both stable and unstable objects in allocentric and egocentric reference frames throughout the task. For egocentric recall, this was at least partially due to the need for participants to “calibrate” their sense of distance within the abstract egocentric coordinate system, whereas the abstract allocentric coordinate system was more intuitive to use. We therefore believe that practice effects are more relevant to egocentric than allocentric learning curves. When examining object-specific learning curves, following previously developed techniques (Gallistel et al., 2004), we observed several differences in allocentric versus egocentric memory, including higher initial performances, higher asymptotic performances, and higher slopes at the change points (i.e., the trials with strongest learning) for allocentric as compared to egocentric memory recall. These effects detail the generally higher level of performance for allocentric compared to egocentric memory recall and suggest that allocentric learning is more abrupt, while egocentric learning is more gradual (though this difference may also be driven by the fact that we did not include objects whose stability was defined egocentrically).

We note that, while the majority of our findings point at distinct determinants of allocentric and egocentric spatial memory, we also found some evidence of mutual influences. At a broad level, we found a positive correlation between allocentric and egocentric memory, indicating that participants with better allocentric memory also excelled in egocentric recall. Furthermore, egocentric memory performance was modulated by the environment’s primary orientation cue, the north boundary. Specifically, egocentric performance was better when the object was close to the north boundary; when participants’ allocentric starting orientation was oriented toward north; and when their allocentric starting orientation was aligned with the major axes of the environment. These effects suggest that participants use some allocentric information for egocentric memory recall, in line with previous observations (Mou and McNamara, 2002; McNamara et al., 2003, 2008; Gagnon et al., 2014). Hence, future studies may investigate the neural underpinnings of how egocentric memory recall involves the use of specific types of allocentric information. When examining the relationship between participants’ demographic characteristics and spatial memory, we observed that older age was associated with poorer allocentric and egocentric memory performance, despite covering only a narrow age range (most participants were between 20 and 30 years old). This is in line with previous reports of a general age-related weakening of spatial memory, irrespective of allocentric and ego-centric reference frames (Colombo et al., 2017). We also found that men performed better on egocentric recall compared to women (with a similar direction of the effect for allocentric recall). This is in line with previous research showing that men outperformed women in both egocentric and allocentric spatial memory tasks (Coutrot et al., 2018; Fernandez-Baizan et al., 2019; Miola et al., 2023). These findings point at interindividual determinants of allocentric and egocentric spatial memory, in addition to the intra-individual determinants discussed above.

Our results may be informative for understanding and modeling the relationship between allocentric and egocentric spatial systems in the human brain. We propose that they are consistent with two coexisting systems that, to some extent, operate with independent streams of input and output and both of which are directly linked to non-spatial object representations (Burgess, 2006). These conclusions follow from our observations of different factors modulating either allocentric or egocentric memory, and the finding that allocentric feedback did not facilitate egocentric performance and *vice versa*. This view does not exclude that there are mutual connections between the two systems at various levels of brain organization, but it speaks against a strictly hierarchical organization of allocentric and egocentric representations, where all sensory information goes through the egocentric system before reaching the allocentric system and where every memory retrieval starts with allocentric representations exclusively linked to non-spatial object information.

Such a hierarchical organization has been suggested, for example, by the Bicanski–Burgess model and its predecessors, which is a computational model that relates various egocentric and allocentric spatial cells to each other and suggests how spatial navigation and memory play out as neural processes that span multiple areas of the brain (Byrne et al., 2007; Bicanski and Burgess, 2018). According to this model, sensory information evokes egocentric representations (in parietal cortices) encoding an individual’s egocentric relationship to objects and boundaries in the environment. This information then travels through a transformation circuit (in the retrosplenial cortex), where it is combined with allocentric head-direction information. This enables the emergence of allocentric representations encoding an individual’s allocentric relationship to objects and boundaries, leading to the emergence of allocentric place representations (in the hippocampus). Non-spatial object information is then attached to these allocentric representations (Bicanski and Burgess, 2018). We believe that such a model implies tighter links between allocentric and egocentric memory performance than the ones we observed. In particular, we believe that the Bicanski–Burgess model would have predicted positive effects of allocentric feedback on egocentric memory performance as the Bicanski–Burgess model allows, in principle, that egocentric relationships to objects are perfectly computable from allocentric representations. In contrast, our results indicate that forgotten egocentric information cannot easily be recovered from allocentric information (otherwise, allocentric feedback should have facilitated egocentric performance; Figure 7). This also implies links between egocentric representations and non-spatial object information that are more direct than proposed before (Bicanski and Burgess, 2018), which may be implemented by synaptic connections between egocentrically tuned neurons and “object-identity cells” at the neural level.

Our results are in line with models of allocentric and egocentric processing streams that are, at least to some extent, parallel to each other (Sholl and Nolin, 1997; Klatzky, 1998; Burgess, 2006; Wang et al., 2020), though these streams may have various mutual interconnections and converge, for example, in the hippocampus. In support of this idea, neuronal recordings in the rat entorhinal cortex revealed that the medial entorhinal cortex is more specialized in processing allocentric information whereas the lateral entorhinal cortex preferentially processes egocentric information (Wang et al., 2018; see also Long et al., 2025 for egocentric tuning in the medial entorhinal cortex). The fact that this allocentric–egocentric distinction occurs in the same structure (the entorhinal cortex) and at a relatively late stage of the processing hierarchy speaks against a strict separation into egocentric representations existing at early stages versus allocentric representations existing at late stages of the hierarchy. Neural recordings during the Garden Game may help further elucidate the extent to which allocentric and egocentric systems are distinct or overlapping, and whether allocentric and egocentric recall recruit the two systems separately or in combination.

The neural processes underlying allocentric and egocentric memory in the Garden Game remain unclear at this stage but they may be supported by various neuronal cell types discovered previously, mostly in rodents (Moser et al., 2017; Bicanski and Burgess, 2020). This includes place cells (O’Keefe and Dostrovsky, 1971; Ekstrom et al., 2003), grid cells (Hafting et al., 2005; Jacobs et al., 2013) and head-direction cells (Taube et al., 1990), which activate when rodents and other animals (potentially including humans) move across particular locations within the environment or along specific global directions. Because the tuning curves of these cells are tied to the external environment, they are considered to be the neural basis of allocentric cognitive maps. Place, grid, and head-direction cells may be active during the encoding periods of the Garden Game, and they may reactivate during allocentric memory recall when participants are asked to remember the locations of objects relative to the environment. Such reinstatement of allocentrically modulated neurons may provide a cellular mechanism for human allocentric spatial memory, but evidence for this idea remains sparse, with only one previous study showing reactivations of human place-cell populations during a free-recall task (Miller et al., 2013).

Conversely, egocentrically tuned neurons that represent directions and distances between the navigating subject and various aspects of the environment may play a critical role for egocentric spatial memory. These neurons may include egocentric boundary-vector cells (Wang et al., 2018; Hinman et al., 2019; Gofman et al., 2019; Alexander et al., 2020), egocentric center-bearing cells (Wang et al., 2018; LaChance et al., 2019), egocentric item-bearing cells (Wang et al., 2018; Jercog et al., 2019; Kunz et al., 2021; Park et al., 2024), and egocentric goal-vector cells (Wilber et al., 2014; Sarel et al., 2017; Wang et al., 2018; Ormond and O’Keefe, 2022). Similar to allocentrically tuned neurons, these cell types may be active during the encoding periods of the Garden Game, and they may reinstate their activity from encoding during egocentric memory recall. A recent study found preliminary evidence for reactivations of egocentrically tuned neurons during human memory recall (Kunz et al., 2021), but future work, for example using the Garden Game, is needed to better understand the exact role of egocentrically tuned neurons in human egocentric spatial memory. This work may also identify to what extent allocentric and egocentric neurons are simultaneously involved in both allocentric and egocentric spatial memory.

Our study has several limitations. First, we used a laptop-based task with a virtual environment, which provides controlled experimental conditions but differs from real-world scenarios (Diersch and Wolbers, 2019; Hejtmanek et al., 2020). Hence, future studies may use virtual-reality setups with head-mounted displays or augmented reality in physically moving participants to create more immersive and realistic settings, helping increase the generalizability of the results to everyday life. Another limitation is the sample size (*n* = 64 in the combined sample), leaving some of the results—such as those related to age and gender—potentially underpowered. Larger studies with the Garden Game may thus reveal additional insights into the determinants of allocentric and egocentric memory. Another limitation is the variability in performance across individual participants and objects. This variability offers interesting opportunities for participant- and object-wise learning analyses (Gallistel et al., 2004), but makes it more challenging to identify general trends. For example, some participants performed very well from the beginning, making it difficult to fit appropriate learning curves with meaningful parameters to their data. We furthermore note that our participants were predominately university students, forming a sample with a relatively narrow age range and limited diversity of educational backgrounds. A broader age and educational range may improve the generalizability of the findings to more diverse populations.

## 5. Conclusion

The Garden Game is an effective tool for investigating human spatial memory. Here, it allowed us to quantify various determinants of allocentric and egocentric spatial memory, suggesting that the underlying systems are at least partially distinct from each other, enabling predictions for future studies with neural recordings. Translational research may use the Garden Game to test for effects of aging and (preclinical) Alzheimer’s disease on spatial memory. This may show whether spatial memory in allocentric or egocentric reference frames is more susceptible to these conditions, as proposed and described before (Moffat, 2009; Harris and Wolbers, 2014; Serino et al., 2014; Tu et al., 2017; Ritchie et al., 2018; Tuena et al., 2021).

## Author contributions statement

Laura Nett: Methodology, Formal analysis, Investigation, Data curation, Writing – Original draft, Visualization, Project administration. Tim A. Guth: Investigation, Writing – Review and editing. Philipp K. Büchel: Writing – Review and editing. Nuttida Rungratsameetaweemana: Conceptualization, Writing – Review and editing. Lukas Kunz: Conceptualization, Methodology, Resources, Software, Writing – Review and editing, Validation, Supervision, Project administration, Funding acquisition.

## Declaration of competing interest

The authors have no competing interests.

## Data and code availability

The data and code used in this project are available in a public GitHub repository, which can be accessed at the following URL: https://github.com/BonnSpatialMemoryLab/ NettGardenGameBehavior2025

## Acknowledgements

We thank the participants for their time and effort. We thank Joshua Jacobs, Florian Mormann, and their research groups for helpful comments on the task design, and we thank Sven Lange for help with data collection.

## Funding

This work was supported by the return program of the Ministry of Culture and Science of the State of North Rhine-Westphalia and by the Federal Ministry of Education and Research (BMBF; 01GQ2402A).

## Notes

https://github.com/BonnSpatialMemoryLab/NettGardenGameBehavior2025

https://github.com/BonnSpatialMemoryLab/GardenGameTask

### Competing Interest Statement

The authors have declared no competing interest.

