## Supplemental File for "Behavioral investigation of allocentric and egocentric cognitive maps in human spatial memory"

Table S1: **Linear mixed models to analyze allocentric memory performance as a function of behavior.**

| Predictor | Cohort | $n$ | $z$ | $p$ |
|---|---|---|---|---|
| **Section 3.2: Associations of time and object stability with memory performance** | | | | |
| Trial index | cohort 1 | 32 | 4.638 | $<0.001^{***}$ |
| Trial index | cohort 2 | 32 | 4.904 | $<0.001^{***}$ |
| Trial index | cohorts 1 and 2 | 64 | 6.748 | $<0.001^{***}$ |
| Trial index (only stable objects) | cohort 1 | 32 | 4.650 | $<0.001^{***}$ |
| Trial index (only stable objects) | cohort 2 | 32 | 3.946 | $<0.001^{***}$ |
| Trial index (only stable objects) | cohorts 1 and 2 | 64 | 6.074 | $<0.001^{***}$ |
| Trial index (only unstable objects) | cohort 1 | 32 | 1.950 | 0.051 |
| Trial index (only unstable objects) | cohort 2 | 32 | 3.000 | $0.003^{*}$ |
| Trial index (only unstable objects) | cohorts 1 and 2 | 64 | 3.498 | $<0.001^{***}$ |
| Object stability (stable > unstable) | cohort 1 | 32 | 3.337 | $0.001^{**}$ |
| Object stability (stable > unstable) | cohort 2 | 32 | 3.017 | $0.003^{**}$ |
| Object stability (stable > unstable) | cohorts 1 and 2 | 64 | 4.492 | $<0.001^{***}$ |
| Trial index : Object stability (stable > unstable) | cohort 1 | 32 | 1.871 | 0.061 |
| Trial index : Object stability (stable > unstable) | cohort 2 | 32 | 0.691 | 0.490 |
| Trial index : Object stability (stable > unstable) | cohorts 1 and 2 | 64 | 1.809 | 0.070 |
| **Section 3.4: Associations of environmental features with memory performance** | | | | |
| Distance to nearest boundary | cohort 1 | 32 | -3.504 | $<0.001^{***}$ |
| Distance to nearest boundary | cohort 2 | 32 | -3.011 | $0.003^{**}$ |
| Distance to nearest boundary | cohorts 1 and 2 | 64 | -4.592 | $<0.001^{***}$ |
| Distance to north boundary | cohort 1 | 32 | -4.260 | $<0.001^{***}$ |
| Distance to north boundary | cohort 2 | 32 | -3.064 | $0.002^{**}$ |
| Distance to north boundary | cohorts 1 and 2 | 64 | -5.166 | $<0.001^{***}$ |
| Distance to nearest corner | cohort 1 | 32 | -4.469 | $<0.001^{***}$ |
| Distance to nearest corner | cohort 2 | 32 | -3.500 | $<0.001^{***}$ |
| Distance to nearest corner | cohorts 1 and 2 | 64 | -5.618 | $<0.001^{***}$ |
| Distance to nearest landmark | cohort 1 | 32 | -0.802 | 0.422 |
| Distance to nearest landmark | cohort 2 | 32 | 1.074 | 0.283 |
| Distance to nearest landmark | cohorts 1 and 2 | 64 | 0.223 | 0.823 |
| **Section 3.5: Associations between starting positions and memory performance** | | | | |
| Distance to starting position | cohort 1 | 32 | -0.507 | 0.612 |
| Distance to starting position | cohort 2 | 32 | -0.867 | 0.386 |
| Distance to starting position | cohorts 1 and 2 | 64 | -0.969 | 0.332 |
| **Section 3.6: Associations between feedback and memory performance** | | | | |
| Allocentric feedback (with > without) | cohort 1 | 32 | 4.816 | $<0.001^{***}$ |
| Allocentric feedback (with > without) | cohort 2 | 32 | 6.265 | $<0.001^{***}$ |
| Allocentric feedback (with > without) | cohorts 1 and 2 | 64 | 7.837 | $<0.001^{***}$ |
| Egocentric feedback (with > without) | cohort 1 | 32 | -2.968 | $0.003^{**}$ |
| Egocentric feedback (with > without) | cohort 2 | 32 | -6.221 | $<0.001^{***}$ |
| Egocentric feedback (with > without) | cohorts 1 and 2 | 64 | -6.498 | $<0.001^{***}$ |
| Retrieval position (2 > 1) | cohort 1 | 32 | 2.209 | $0.027^{*}$ |
| Retrieval position (2 > 1) | cohort 2 | 32 | 3.181 | $0.001^{**}$ |
| Retrieval position (2 > 1) | cohorts 1 and 2 | 64 | 3.810 | $<0.001^{***}$ |
| Retrieval position (3 > 1) | cohort 1 | 32 | -3.307 | $0.001^{**}$ |
| Retrieval position (3 > 1) | cohort 2 | 32 | -5.695 | $<0.001^{***}$ |
| Retrieval position (3 > 1) | cohorts 1 and 2 | 64 | -6.363 | $<0.001^{***}$ |
| Retrieval position (4 > 1) | cohort 1 | 32 | 1.304 | 0.192 |
| Retrieval position (4 > 1) | cohort 2 | 32 | 0.031 | 0.976 |
| Retrieval position (4 > 1) | cohorts 1 and 2 | 64 | 0.944 | 0.345 |
| Time since end of encoding | cohort 1 | 32 | -2.428 | $0.015^{*}$ |
| Time since end of encoding | cohort 2 | 32 | -4.226 | $<0.001^{***}$ |

*Continued on next page*

| Predictor | Cohort | $n$ | $z$ | $p$ |
|---|---|---|---|---|
| Time since end of encoding | cohorts 1 and 2 | 64 | -4.659 | < 0.001*** |
| Time since encoding of firstly encoded object (only firstly encoded object) | cohort 1 | 32 | -2.033 | 0.042* |
| Time since encoding of firstly encoded object (only firstly encoded object) | cohort 2 | 32 | -1.843 | 0.065 |
| Time since encoding of firstly encoded object (only firstly encoded object) | cohorts 1 and 2 | 64 | -2.721 | 0.007** |
| Time since encoding of secondly encoded object (only secondly encoded object) | cohort 1 | 32 | -1.915 | 0.055 |
| Time since encoding of secondly encoded object (only secondly encoded object) | cohort 2 | 32 | -3.348 | 0.001** |
| Time since encoding of secondly encoded object (only secondly encoded object) | cohorts 1 and 2 | 64 | -3.692 | < 0.001*** |
| **Section 3.7: Associations between viewing behavior and memory performance** | | | | |
| Time viewing the boundaries (relative to encoding) | cohort 2 | 32 | 2.473 | 0.013* |
| Time viewing the north boundary (relative to encoding) | cohort 2 | 32 | 6.663 | < 0.001*** |
| Time viewing the east boundary (relative to encoding) | cohort 2 | 32 | -0.870 | 0.384 |
| Time viewing the south boundary (relative to encoding) | cohort 2 | 32 | -3.445 | 0.001** |
| Time viewing the west boundary (relative to encoding) | cohort 2 | 32 | -0.163 | 0.870 |
| Time viewing the corners (relative to encoding) | cohort 2 | 32 | 3.084 | 0.002** |
| Time viewing the north east corner (relative to encoding) | cohort 2 | 32 | 2.690 | 0.007** |
| Time viewing the south east corner (relative to encoding) | cohort 2 | 32 | -0.844 | 0.398 |
| Time viewing the south west corner (relative to encoding) | cohort 2 | 32 | -1.355 | 0.175 |
| Time viewing the north west corner (relative to encoding) | cohort 2 | 32 | 3.407 | 0.001** |
| Time viewing the object (relative to encoding) | cohort 2 | 32 | -0.875 | 0.381 |
| Time viewing the gaze area (relative to encoding) | cohort 2 | 32 | 1.539 | 0.124 |
| Time viewing the object and gaze area (relative to encoding) | cohort 2 | 32 | -0.714 | 0.475 |
| Gaze coverage | cohort 2 | 32 | -1.330 | 0.183 |
| **Section 3.8: Age and gender effects** | | | | |
| Age *(controlling for gender)* | cohort 1 | 32 | 1.174 | 0.240 |
| Age *(controlling for gender)* | cohort 2 | 32 | -2.321 | 0.020* |
| Age *(controlling for gender)* | cohorts 1 and 2 | 64 | -1.570 | 0.116 |
| Age *(controlling for gender)* | complete dataset | 80 | -2.241 | 0.025* |
| Gender (male > female) *(controlling for age)* | cohort 1 | 32 | -1.703 | 0.089 |
| Gender (male > female) *(controlling for age)* | cohort 2 | 32 | 2.325 | 0.020* |
| Gender (male > female) *(controlling for age)* | cohorts 1 and 2 | 64 | 0.932 | 0.351 |
| Gender (male > female) *(controlling for age)* | complete dataset | 80 | 1.067 | 0.286 |

Cohorts 1 and 2 contain 32 participants each. The complete dataset additionally includes $n = 16$ participants who completed a slightly different version of the Garden Game. $^{*}p < 0.05$; $^{**}p < 0.01$; $^{***}p < 0.001$.

Table S2: **Linear mixed models to analyze egocentric memory performance as a function of behavior.**

| **Predictor** | **Cohort** | ***n*** | ***z*** | ***p*** |
|---|---|---|---|---|
| **Section 3.2: Association of time and object stability with memory performance** | | | | |
| Trial index | cohort 1 | 32 | 6.775 | $<0.001^{***}$ |
| Trial index | cohort 2 | 32 | 6.867 | $<0.001^{***}$ |
| Trial index | cohorts 1 and 2 | 64 | 9.629 | $<0.001^{***}$ |
| Trial index (only stable objects) | cohort 1 | 32 | 4.516 | $<0.001^{***}$ |
| Trial index (only stable objects) | cohort 2 | 32 | 6.171 | $<0.001^{***}$ |
| Trial index (only stable objects) | cohorts 1 and 2 | 64 | 7.542 | $<0.001^{***}$ |
| Trial index (only unstable objects) | cohort 1 | 32 | 5.008 | $<0.001^{***}$ |
| Trial index (only unstable objects) | cohort 2 | 32 | 3.507 | $<0.001^{***}$ |
| Trial index (only unstable objects) | cohorts 1 and 2 | 64 | 6.049 | $<0.001^{***}$ |
| Object stability (stable > unstable) | cohort 1 | 32 | -0.213 | 0.832 |
| Object stability (stable > unstable) | cohort 2 | 32 | -0.508 | 0.611 |
| Object stability (stable > unstable) | cohorts 1 and 2 | 64 | -0.505 | 0.613 |
| Trial index : Object stability (stable > unstable) | cohort 1 | 32 | -0.410 | 0.682 |
| Trial index : Object stability (stable > unstable) | cohort 2 | 32 | 1.932 | 0.053 |
| Trial index : Object stability (stable > unstable) | cohorts 1 and 2 | 64 | 1.042 | 0.297 |
| **Section 3.4: Associations of environmental features with memory performance** | | | | |
| Distance to nearest boundary | cohort 1 | 32 | 1.926 | 0.054 |
| Distance to nearest boundary | cohort 2 | 32 | 1.336 | 0.181 |
| Distance to nearest boundary | cohorts 1 and 2 | 64 | 2.299 | $0.022^{*}$ |
| Distance to nearest boundary *(controlling for distance to starting position)* | cohort 1 | 32 | 0.962 | 0.336 |
| Distance to nearest boundary *(controlling for distance to starting position)* | cohort 2 | 32 | -0.235 | 0.814 |
| Distance to nearest boundary *(controlling for distance to starting position)* | cohorts 1 and 2 | 64 | 0.531 | 0.595 |
| Distance to north boundary | cohort 1 | 32 | -3.393 | $0.001^{**}$ |
| Distance to north boundary | cohort 2 | 32 | -2.174 | $0.030^{*}$ |
| Distance to north boundary | cohorts 1 and 2 | 64 | -3.940 | $<0.001^{***}$ |
| Distance to north boundary *(controlling for distance to starting position)* | cohort 1 | 32 | -3.308 | $0.001^{**}$ |
| Distance to north boundary *(controlling for distance to starting position)* | cohort 2 | 32 | -2.423 | $0.015^{*}$ |
| Distance to north boundary *(controlling for distance to starting position)* | cohorts 1 and 2 | 64 | -4.032 | $<0.001^{***}$ |
| Distance to nearest corner | cohort 1 | 32 | 2.155 | $0.031^{*}$ |
| Distance to nearest corner | cohort 2 | 32 | 1.415 | 0.157 |
| Distance to nearest corner | cohorts 1 and 2 | 64 | 2.517 | $0.012^{*}$ |
| Distance to nearest corner *(controlling for distance to starting position)* | cohort 1 | 32 | 1.218 | 0.223 |
| Distance to nearest corner *(controlling for distance to starting position)* | cohort 2 | 32 | -0.072 | 0.943 |
| Distance to nearest corner *(controlling for distance to starting position)* | cohorts 1 and 2 | 64 | 0.824 | 0.410 |
| Distance to nearest landmark | cohort 1 | 32 | -1.159 | 0.247 |
| Distance to nearest landmark | cohort 2 | 32 | 1.466 | 0.143 |
| Distance to nearest landmark | cohorts 1 and 2 | 64 | 0.238 | 0.812 |
| **Section 3.5: Associations between starting positions and memory performance** | | | | |
| Distance to starting position | cohort 1 | 32 | -4.077 | $<0.001^{***}$ |
| Distance to starting position | cohort 2 | 32 | -5.296 | $<0.001^{***}$ |
| Distance to starting position | cohorts 1 and 2 | 64 | -6.615 | $<0.001^{***}$ |

*Continued on next page*

| **Predictor** | **Cohort** | ***n*** | ***z*** | ***p*** |
|---|---|---|---|---|
| Alignment with the cardinal axes (aligned > misaligned) | cohort 1 | 32 | 3.671 | < 0.001*** |
| Alignment with the cardinal axes (aligned > misaligned) | cohort 2 | 32 | 4.948 | < 0.001*** |
| Alignment with the cardinal axes (aligned > misaligned) | cohorts 1 and 2 | 64 | 6.075 | < 0.001*** |
| **Section 3.6: Associations between feedback and memory performance** | | | | |
| Allocentric feedback (with > without) | cohort 1 | 32 | -3.379 | 0.001** |
| Allocentric feedback (with > without) | cohort 2 | 32 | -2.252 | 0.024* |
| Allocentric feedback (with > without) | cohorts 1 and 2 | 64 | -3.997 | < 0.001*** |
| Egocentric feedback (with > without) | cohort 1 | 32 | -1.327 | 0.184 |
| Egocentric feedback (with > without) | cohort 2 | 32 | -1.192 | 0.233 |
| Egocentric feedback (with > without) | cohorts 1 and 2 | 64 | -1.783 | 0.075 |
| Retrieval position (2 > 1) | cohort 1 | 32 | -1.587 | 0.113 |
| Retrieval position (2 > 1) | cohort 2 | 32 | -1.190 | 0.234 |
| Retrieval position (2 > 1) | cohorts 1 and 2 | 64 | -1.969 | 0.049* |
| Retrieval position (3 > 1) | cohort 1 | 32 | -3.037 | 0.002** |
| Retrieval position (3 > 1) | cohort 2 | 32 | -1.939 | 0.052 |
| Retrieval position (3 > 1) | cohorts 1 and 2 | 64 | -3.533 | < 0.001*** |
| Retrieval position (4 > 1) | cohort 1 | 32 | -3.330 | 0.001** |
| Retrieval position (4 > 1) | cohort 2 | 32 | -2.436 | 0.015* |
| Retrieval position (4 > 1) | cohorts 1 and 2 | 64 | -4.089 | < 0.001*** |
| Time since end of encoding | cohort 1 | 32 | -6.234 | < 0.001*** |
| Time since end of encoding | cohort 2 | 32 | -3.668 | < 0.001*** |
| Time since end of encoding | cohorts 1 and 2 | 64 | -7.107 | < 0.001*** |
| Time since encoding of first encoded object (only first encoded object) | cohort 1 | 32 | -3.054 | 0.002** |
| Time since encoding of first encoded object (only first encoded object) | cohort 2 | 32 | -2.469 | 0.014* |
| Time since encoding of first encoded object (only first encoded object) | cohorts 1 and 2 | 64 | -3.935 | < 0.001*** |
| Time since encoding of second encoded object (only first encoded object) | cohort 1 | 32 | -5.897 | < 0.001*** |
| Time since encoding of second encoded object (only second encoded object) | cohort 2 | 32 | -3.180 | 0.001** |
| Time since encoding of second encoded object (only second encoded object) | cohorts 1 and 2 | 64 | -6.541 | < 0.001*** |
| **Section 3.7: Associations between viewing behavior and memory performance** | | | | |
| Time viewing the boundaries (relative to encoding) | cohort 2 | 32 | -2.657 | 0.008** |
| Time viewing the north boundary (relative to encoding) | cohort 2 | 32 | 1.189 | 0.234 |
| Time viewing the east boundary (relative to encoding) | cohort 2 | 32 | -1.147 | 0.251 |
| Time viewing the south boundary (relative to encoding) | cohort 2 | 32 | -3.163 | 0.002** |
| Time viewing the west boundary (relative to encoding) | cohort 2 | 32 | -0.179 | 0.858 |
| Time viewing the corners (relative to encoding) | cohort 2 | 32 | 0.167 | 0.868 |
| Time viewing the north east corner (relative to encoding) | cohort 2 | 32 | 0.396 | 0.692 |
| Time viewing the south east corner (relative to encoding) | cohort 2 | 32 | -2.045 | 0.041* |

*Continued on next page*

| **Predictor** | **Cohort** | $n$ | $z$ | $p$ |
|---|---|---|---|---|
| Time viewing the south west corner (relative to encoding) | cohort 2 | 32 | -0.083 | 0.934 |
| Time viewing the north west corner (relative to encoding) | cohort 2 | 32 | 1.553 | 0.120 |
| Time viewing the object (relative to encoding) | cohort 2 | 32 | 1.859 | 0.063 |
| Time viewing the gaze area (relative to encoding) | cohort 2 | 32 | 2.034 | 0.042* |
| Time viewing the object and gaze area (relative to encoding) | cohort 2 | 32 | 2.053 | 0.040* |
| Gaze coverage | cohort 2 | 32 | -4.523 | < 0.001*** |
| **Section 3.8 Age and Gender Effects** | | | | |
| Age *(controlling for gender)* | cohort 1 | 32 | -0.449 | 0.653 |
| Age *(controlling for gender)* | cohort 2 | 32 | -1.650 | 0.099 |
| Age *(controlling for gender)* | cohorts 1 and 2 | 64 | -1.743 | 0.081 |
| Age *(controlling for gender)* | complete dataset | 80 | -2.126 | 0.034* |
| Gender (male > female) *(controlling for age)* | cohort 1 | 32 | 0.737 | 0.461 |
| Gender (male > female) *(controlling for age)* | cohort 2 | 32 | 2.528 | 0.011* |
| Gender (male > female) *(controlling for age)* | cohorts 1 and 2 | 64 | 2.445 | 0.014* |
| Gender (male > female) *(controlling for age)* | complete dataset | 80 | 2.411 | 0.016* |

Cohorts 1 and 2 contain 32 participants each. The complete dataset additionally includes $n = 16$ participants who completed a slightly different version of the Garden Game. $^{*}p < 0.05$; $^{**}p < 0.01$; $^{***}p < 0.001$.

Table S3: **Statistical information for analyses in the main text, excluding linear mixed models.**

| **Test** | **Statistics** |
|---|---|
| **Section 2.3: Participants** | |
| Total number of subjects in cohort 1 | $n = 32$. |
| Total number of subjects in cohort 2 | $n = 32$. |
| Total number of additional subjects | $n = 16$. |
| Average age in cohort 1 | $25 \pm 3$ (mean ± SD); range, 18 – 33. |
| Average age in cohort 2 | $25 \pm 4$ (mean ± SD); range, 20 – 40. |
| Average age in the additional subjects | $27 \pm 7$ (mean ± SD); range, 19 – 53. |
| Number of females in cohort 1 | $n_{female} = 17$. |
| Number of females in cohort 2 | $n_{female} = 17$. |
| Number of females in the additional subjects | $n_{female} = 7$. |
| **Section 3.1 Behavioral task.** | |
| Duration per session (minutes) | $66.19 \pm 1.56$ (mean ± SEM); range, 42.85 – 96.39. |
| Duration of starting position per session (minutes) | $5.06 \pm 0.20$ (mean ± SEM); range, 2.94 – 9.31. |
| Duration of encoding per session (minutes) | $25.22 \pm 0.84$ (mean ± SEM); range, 13.01 – 43.42. |
| Duration of allocentric retrieval per session (minutes) | $9.33 \pm 0.36$ (mean ± SEM); range, 4.61 – 20.02. |
| Duration of egocentric retrieval per session (minutes) | $12.25 \pm 0.60$ (mean ± SEM); range, 5.67 – 28.58. |
| Duration of score screen per session (minutes) | $1.59 \pm 0.14$ (mean ± SEM); range, 0.38 – 5.88. |
| **Section 3.2: Associations of time and object stability with memory performance** | |
| Paired $t$-test between allocentric and egocentric memory performance | $t = 5.250$, $p < 0.001^{***}$, $n = 32$ (cohort 1)<br>$t = 7.683$, $p < 0.001^{***}$, $n = 32$ (cohort 2)<br>$t = 8.566$, $p < 0.001^{***}$, $n = 64$ (cohorts 1 and 2) |
| Pearson correlation between allocentric and egocentric memory performance | $r = 0.536$, $p = 0.002^{**}$, $n = 32$ (cohort 1)<br>$r = 0.878$, $p < 0.001^{***}$, $n = 32$ (cohort 2)<br>$r = 0.734$, $p < 0.001^{***}$, $n = 64$ (cohorts 1 and 2) |
| LMM with performance as the dependent variable and an interaction between retrieval type (egocentric > allocentric) and object stability (stable > unstable) | $z = -2.385$, $p = 0.017^{*}$, $n = 32$ (cohort 1)<br>$z = -2.451$, $p = 0.014^{*}$, $n = 32$ (cohort 2)<br>$z = -3.419$, $p = 0.001^{**}$, $n = 64$ (cohorts 1 and 2) |
| **Section 3.3: Object-specific learning** | |
| Total number of change points in allocentric learning curves | $n = 108$ (of 384). |
| Total number of change points in egocentric learning curves | $n = 151$ (of 384). |
| Total number of change points for stable objects regarding allocentric learning curves | $n = 58$ (of 108). |
| Total number of change points for unstable objects regarding allocentric learning curves | $n = 50$ (of 108). |
| Total number of change points for stable objects regarding egocentric learning curves | $n = 78$ (of 151). |
| Total number of change points for unstable objects regarding egocentric learning curves | $n = 73$ (of 151). |
| $\chi^2$-test between the number of change points for egocentric vs. allocentric learning curves | $\chi^2 = 10.276$, $p = 0.001^{**}$, $n = 384$. |
| $\chi^2$-test between the number of change points for stable vs. unstable objects regarding allocentric learning curves | $\chi^2 = 0.631$, $p = 0.427$, $n = 192$. |

*Continued on next page*

| Test | Statistics |
| --- | --- |
| $\chi^2$-test between the number of change points for stable vs. unstable objects regarding egocentric learning curves | $\chi^2 = 0.175$, $p = 0.676$, $n = 192$. |
| t-test between the initial performance for allocentric vs. egocentric learning curves | $t = 6.864$, $p < 0.001^{***}$, $n = 384$. |
| t-test between the initial performance for stable vs. unstable objects regarding allocentric learning curves | $t = -0.167$, $p = 0.867$, $n = 192$. |
| t-test between the initial performance for stable vs. unstable objects regarding egocentric learning curves | $t = -0.913$, $p = 0.362$, $n = 192$. |
| t-test between the asymptotic performance for allocentric vs. egocentric learning curves | $t = 9.518$, $p < 0.001^{***}$, $n = 384$. |
| t-test between the asymptotic performance for stable vs. unstable objects regarding allocentric learning curves | $t = 2.836$, $p = 0.005^{**}$, $n = 192$. |
| t-test between the asymptotic performance for stable vs. unstable objects regarding egocentric learning curves | $t = 0.095$, $p = 0.925$, $n = 192$. |
| t-test between the trial index with the change point for allocentric vs. egocentric learning curves | $t = -0.295$, $p = 0.768$, $n_{allo} = 108, n_{ego} = 151$. |
| t-test between the trial index with the change point for stable vs. unstable objects regarding allocentric learning curves | $t = 1.344$, $p = 0.182$, $n_{stable} = 58, n_{unstable} = 50$. |
| t-test between the trial index with the change point for stable vs. unstable objects regarding egocentric learning curves | $t = 0.825$, $p = 0.411$, $n_{stable} = 78, n_{unstable} = 73$. |
| t-test between the slope at the change point for allocentric vs. egocentric learning curves | $t = 3.128$, $p = 0.002^{**}$, $n_{allo} = 108, n_{ego} = 151$. |
| t-test between the slope at the change point for stable vs. unstable objects regarding allocentric learning curves | $t = -1.156$, $p = 0.251$, $n_{stable} = 58, n_{unstable} = 50$. |
| t-test between the slope at the change point for stable vs. unstable objects regarding egocentric learning curves | $t = 0.123$, $p = 0.902$, $n_{stable} = 78, n_{unstable} = 73$. |
| **Section 3.5: Associations between starting positions and memory performance** | |
| Friedman-test to examine the relationship between allocentric starting orientation and allocentric memory performance | $F = 5.192$, $p < 0.001^{***}$, $n = 32$ (cohort 1)<br>$F = 7.750$, $p < 0.001^{***}$, $n = 32$ (cohort 2)<br>$F = 11.360$, $p < 0.001^{***}$, $n = 64$ (cohorts 1 and 2) |
| Friedman-test to examine the relationship between allocentric starting orientation and egocentric memory performance | $F = 3.869$, $p < 0.001^{***}$, $n = 32$ (cohort 1)<br>$F = 6.329$, $p < 0.001^{***}$, $n = 32$ (cohort 2)<br>$F = 9.255$, $p < 0.001^{***}$, $n = 64$ (cohorts 1 and 2) |
| Friedman-test to examine the relationship between egocentric starting orientation and allocentric memory performance | $F = 1.247$, $p = 0.255$, $n = 32$ (cohort 1)<br>$F = 0.248$, $p = 0.994$, $n = 32$ (cohort 2)<br>$F = 0.736$, $p = 0.704$, $n = 64$ (cohorts 1 and 2) |
| Friedman-test to examine the relationship between egocentric starting orientation and egocentric memory performance | $F = 5.464$, $p < 0.001^{***}$, $n = 32$ (cohort 1)<br>$F = 7.133$, $p < 0.001^{***}$, $n = 32$ (cohort 2)<br>$F = 11.396$, $p < 0.001^{***}$, $n = 64$ (cohorts 1 and 2) |

$^{*}p < 0.05$; $^{**}p < 0.01$; $^{***}p < 0.001$.

Table S4: **Results of Wilcoxon signed rank test with Sidak correction for multiple comparisons to examine the relationship between egocentric starting orientation and egocentric memory performance.**

| **Cohort** | $n$ | **Orientation 1** | **Orientation 2** | $W$ | $p_{corr.}$ |
|---|---|---|---|---|---|
| Cohort 1 | 32 | A | AR | 216 | 1.000 |
| Cohort 2 | 32 | A | AR | 119 | 0.315 |
| Cohorts 1 and 2 | 64 | A | AR | 632 | 0.344 |
| Cohort 1 | 32 | A | RA | 135 | 0.626 |
| Cohort 2 | 32 | A | RA | 73 | 0.011* |
| Cohorts 1 and 2 | 64 | A | RA | 406 | 0.002** |
| Cohort 1 | 32 | A | R | 92 | 0.054 |
| Cohort 2 | 32 | A | R | 74 | 0.012* |
| Cohorts 1 and 2 | 64 | A | R | 330 | <0.001*** |
| Cohort 1 | 32 | A | RB | 68 | 0.007* |
| Cohort 2 | 32 | A | RB | 22 | <0.001*** |
| Cohorts 1 and 2 | 64 | A | RB | 156 | <0.001*** |
| Cohort 1 | 32 | A | BR | 63 | 0.004* |
| Cohort 2 | 32 | A | BR | 36 | <0.001*** |
| Cohorts 1 and 2 | 64 | A | BR | 193 | <0.001*** |
| Cohort 1 | 32 | A | B | 74 | 0.012* |
| Cohort 2 | 32 | A | B | 58 | 0.002* |
| Cohorts 1 and 2 | 64 | A | B | 254 | <0.001*** |
| Cohort 1 | 32 | A | BL | 65 | 0.005* |
| Cohort 2 | 32 | A | BL | 23 | <0.001*** |
| Cohorts 1 and 2 | 64 | A | BL | 161 | <0.001*** |
| Cohort 1 | 32 | A | LB | 36 | <0.001*** |
| Cohort 2 | 32 | A | LB | 70 | 0.008* |
| Cohorts 1 and 2 | 64 | A | LB | 206 | <0.001*** |
| Cohort 1 | 32 | A | L | 49 | <0.001*** |
| Cohort 2 | 32 | A | L | 56 | 0.002** |
| Cohorts 1 and 2 | 64 | A | L | 201 | <0.001*** |
| Cohort 1 | 32 | A | LA | 81 | 0.022* |
| Cohort 2 | 32 | A | LA | 54 | 0.002* |
| Cohorts 1 and 2 | 64 | A | LA | 262 | <0.001*** |
| Cohort 1 | 32 | A | AL | 166 | 0.990 |
| Cohort 2 | 32 | A | AL | 110 | 0.188 |
| Cohorts 1 and 2 | 64 | A | AL | 516 | 0.030* |
| Cohort 1 | 32 | AR | RA | 178 | 1.000 |
| Cohort 2 | 32 | AR | RA | 203 | 1.000 |
| Cohorts 1 and 2 | 64 | AR | RA | 745 | 0.963 |
| Cohort 1 | 32 | AR | R | 102 | 0.112 |
| Cohort 2 | 32 | AR | R | 184 | 1.000 |
| Cohorts 1 and 2 | 64 | AR | R | 546 | 0.061 |
| Cohort 1 | 32 | AR | RB | 103 | 0.120 |
| Cohort 2 | 32 | AR | RB | 68 | 0.007* |
| Cohorts 1 and 2 | 64 | AR | RB | 332 | <0.001*** |
| Cohort 1 | 32 | AR | BR | 111 | 0.200 |
| Cohort 2 | 32 | AR | BR | 69 | 0.007* |
| Cohorts 1 and 2 | 64 | AR | BR | 354 | <0.001*** |
| Cohort 1 | 32 | AR | B | 111 | 0.223 |
| Cohort 2 | 32 | AR | B | 180 | 1.000 |
| Cohorts 1 and 2 | 64 | AR | B | 539 | 0.052 |
| Cohort 1 | 32 | AR | BL | 131 | 0.543 |
| Cohort 2 | 32 | AR | BL | 129 | 0.502 |
| Cohorts 1 and 2 | 64 | AR | BL | 507 | 0.024* |

*Continued on next page*

| Cohort | $n$ | Orientation 1 | Orientation 2 | $W$ | $p_{corr.}$ |
|---|---|---|---|---|---|
| Cohort 1 | 32 | AR | LB | 71 | 0.009* |
| Cohort 2 | 32 | AR | LB | 139 | 0.707 |
| Cohorts 1 and 2 | 64 | AR | LB | 403 | 0.001* |
| Cohort 1 | 32 | AR | L | 112 | 0.213 |
| Cohort 2 | 32 | AR | L | 156 | 0.946 |
| Cohorts 1 and 2 | 64 | AR | L | 506 | 0.023* |
| Cohort 1 | 32 | AR | LA | 107 | 0.156 |
| Cohort 2 | 32 | AR | LA | 171 | 0.997 |
| Cohorts 1 and 2 | 64 | AR | LA | 527 | 0.039* |
| Cohort 1 | 32 | AR | AL | 235 | 1.000 |
| Cohort 2 | 32 | AR | AL | 254 | 1.000 |
| Cohorts 1 and 2 | 64 | AR | AL | 964 | 1.000 |
| Cohort 1 | 32 | RA | R | 142 | 0.764 |
| Cohort 2 | 32 | RA | R | 231 | 1.000 |
| Cohorts 1 and 2 | 64 | RA | R | 761 | 0.985 |
| Cohort 1 | 32 | RA | RB | 183 | 1.000 |
| Cohort 2 | 32 | RA | RB | 104 | 0.128 |
| Cohorts 1 and 2 | 64 | RA | RB | 547 | 0.063 |
| Cohort 1 | 32 | RA | BR | 199 | 1.000 |
| Cohort 2 | 32 | RA | BR | 98 | 0.084 |
| Cohorts 1 and 2 | 64 | RA | BR | 602 | 0.201 |
| Cohort 1 | 32 | RA | B | 193 | 1.000 |
| Cohort 2 | 32 | RA | B | 224 | 1.000 |
| Cohorts 1 and 2 | 64 | RA | B | 820 | 1.000 |
| Cohort 1 | 32 | RA | BL | 233 | 1.000 |
| Cohort 2 | 32 | RA | BL | 167 | 0.993 |
| Cohorts 1 and 2 | 64 | RA | BL | 777 | 0.996 |
| Cohort 1 | 32 | RA | LB | 114 | 0.239 |
| Cohort 2 | 32 | RA | LB | 170 | 0.996 |
| Cohorts 1 and 2 | 64 | RA | LB | 562 | 0.088 |
| Cohort 1 | 32 | RA | L | 146 | 0.831 |
| Cohort 2 | 32 | RA | L | 214 | 1.000 |
| Cohorts 1 and 2 | 64 | RA | L | 707 | 0.824 |
| Cohort 1 | 32 | RA | LA | 185 | 1.000 |
| Cohort 2 | 32 | RA | LA | 224 | 1.000 |
| Cohorts 1 and 2 | 64 | RA | LA | 797 | 0.999 |
| Cohort 1 | 32 | RA | AL | 204 | 1.000 |
| Cohort 2 | 32 | RA | AL | 225 | 1.000 |
| Cohorts 1 and 2 | 64 | RA | AL | 837 | 1.000 |
| Cohort 1 | 32 | R | RB | 225 | 1.000 |
| Cohort 2 | 32 | R | RB | 78 | 0.017* |
| Cohorts 1 and 2 | 64 | R | RB | 611 | 0.238 |
| Cohort 1 | 32 | R | BR | 241 | 1.000 |
| Cohort 2 | 32 | R | BR | 158 | 0.960 |
| Cohorts 1 and 2 | 64 | R | BR | 767 | 0.990 |
| Cohort 1 | 32 | R | B | 215 | 1.000 |
| Cohort 2 | 32 | R | B | 223 | 1.000 |
| Cohorts 1 and 2 | 64 | R | B | 875 | 1.000 |
| Cohort 1 | 32 | R | BL | 262 | 1.000 |
| Cohort 2 | 32 | R | BL | 176 | 0.999 |
| Cohorts 1 and 2 | 64 | R | BL | 874 | 1.000 |
| Cohort 1 | 32 | R | LB | 226 | 1.000 |
| Cohort 2 | 32 | R | LB | 206 | 1.000 |
| Cohorts 1 and 2 | 64 | R | LB | 845 | 1.000 |
| Cohort 1 | 32 | R | L | 200 | 1.000 |

*Continued on next page*

| Cohort | $n$ | Orientation 1 | Orientation 2 | $W$ | $p_{corr.}$ |
|---|---|---|---|---|---|
| Cohort 2 | 32 | R | L | 198 | 1.000 |
| Cohorts 1 and 2 | 64 | R | L | 778 | 0.996 |
| Cohort 1 | 32 | R | LA | 213 | 1.000 |
| Cohort 2 | 32 | R | LA | 198 | 1.000 |
| Cohorts 1 and 2 | 64 | R | LA | 818 | 1.000 |
| Cohort 1 | 32 | R | AL | 153 | 0.920 |
| Cohort 2 | 32 | R | AL | 244 | 1.000 |
| Cohorts 1 and 2 | 64 | R | AL | 759 | 0.983 |
| Cohort 1 | 32 | RB | BR | 239 | 1.000 |
| Cohort 2 | 32 | RB | BR | 210 | 1.000 |
| Cohorts 1 and 2 | 64 | RB | BR | 892 | 1.000 |
| Cohort 1 | 32 | RB | B | 247 | 1.000 |
| Cohort 2 | 32 | RB | B | 125 | 0.423 |
| Cohorts 1 and 2 | 64 | RB | B | 729 | 0.920 |
| Cohort 1 | 32 | RB | BL | 218 | 1.000 |
| Cohort 2 | 32 | RB | BL | 174 | 0.999 |
| Cohorts 1 and 2 | 64 | RB | BL | 768 | 0.991 |
| Cohort 1 | 32 | RB | LB | 228 | 1.000 |
| Cohort 2 | 32 | RB | LB | 171 | 0.997 |
| Cohorts 1 and 2 | 64 | RB | LB | 939 | 1.000 |
| Cohort 1 | 32 | RB | L | 222 | 1.000 |
| Cohort 2 | 32 | RB | L | 154 | 0.930 |
| Cohorts 1 and 2 | 64 | RB | L | 916 | 1.000 |
| Cohort 1 | 32 | RB | LA | 246 | 1.000 |
| Cohort 2 | 32 | RB | LA | 158 | 0.960 |
| Cohorts 1 and 2 | 64 | RB | LA | 798 | 0.999 |
| Cohort 1 | 32 | RB | AL | 135 | 0.626 |
| Cohort 2 | 32 | RB | AL | 56 | 0.002* |
| Cohorts 1 and 2 | 64 | RB | AL | 376 | $<$0.001*** |
| Cohort 1 | 32 | BR | B | 253 | 1.000 |
| Cohort 2 | 32 | BR | B | 150 | 0.887 |
| Cohorts 1 and 2 | 64 | BR | B | 798 | 0.999 |
| Cohort 1 | 32 | BR | BL | 192 | 1.000 |
| Cohort 2 | 32 | BR | BL | 208 | 1.000 |
| Cohorts 1 and 2 | 64 | BR | BL | 797 | 0.999 |
| Cohort 1 | 32 | BR | LB | 203 | 1.000 |
| Cohort 2 | 32 | BR | LB | 184 | 1.000 |
| Cohorts 1 and 2 | 64 | BR | LB | 1004 | 1.000 |
| Cohort 1 | 32 | BR | L | 207 | 1.000 |
| Cohort 2 | 32 | BR | L | 184 | 1.000 |
| Cohorts 1 and 2 | 64 | BR | L | 997 | 1.000 |
| Cohort 1 | 32 | BR | LA | 261 | 1.000 |
| Cohort 2 | 32 | BR | LA | 184 | 1.000 |
| Cohorts 1 and 2 | 64 | BR | LA | 893 | 1.000 |
| Cohort 1 | 32 | BR | AL | 147 | 0.846 |
| Cohort 2 | 32 | BR | AL | 74 | 0.012* |
| Cohorts 1 and 2 | 64 | BR | AL | 434 | 0.003* |
| Cohort 1 | 32 | B | BL | 225 | 1.000 |
| Cohort 2 | 32 | B | BL | 204 | 1.000 |
| Cohorts 1 and 2 | 64 | B | BL | 981 | 1.000 |
| Cohort 1 | 32 | B | LB | 216 | 1.000 |
| Cohort 2 | 32 | B | LB | 260 | 1.000 |
| Cohorts 1 and 2 | 64 | B | LB | 937 | 1.000 |
| Cohort 1 | 32 | B | L | 227 | 1.000 |
| Cohort 2 | 32 | B | L | 260 | 1.000 |

*Continued on next page*

| **Cohort** | $n$ | **Orientation 1** | **Orientation 2** | $W$ | $p_{corr.}$ |
|---|---|---|---|---|---|
| Cohorts 1 and 2 | 64 | B | L | 951 | 1.000 |
| Cohort 1 | 32 | B | LA | 258 | 1.000 |
| Cohort 2 | 32 | B | LA | 256 | 1.000 |
| Cohorts 1 and 2 | 64 | B | LA | 1027 | 1.000 |
| Cohort 1 | 32 | B | AL | 118 | 0.299 |
| Cohort 2 | 32 | B | AL | 172 | 0.998 |
| Cohorts 1 and 2 | 64 | B | AL | 570 | 0.105 |
| Cohort 1 | 32 | BL | LB | 171 | 0.997 |
| Cohort 2 | 32 | BL | LB | 220 | 1.000 |
| Cohorts 1 and 2 | 64 | BL | LB | 937 | 1.000 |
| Cohort 1 | 32 | BL | L | 161 | 0.975 |
| Cohort 2 | 32 | BL | L | 224 | 1.000 |
| Cohorts 1 and 2 | 64 | BL | L | 929 | 1.000 |
| Cohort 1 | 32 | BL | LA | 228 | 1.000 |
| Cohort 2 | 32 | BL | LA | 224 | 1.000 |
| Cohorts 1 and 2 | 64 | BL | LA | 1018 | 1.000 |
| Cohort 1 | 32 | BL | AL | 158 | 0.960 |
| Cohort 2 | 32 | BL | AL | 135 | 0.626 |
| Cohorts 1 and 2 | 64 | BL | AL | 575 | 0.116 |
| Cohort 1 | 32 | LB | L | 251 | 1.000 |
| Cohort 2 | 32 | LB | L | 241 | 1.000 |
| Cohorts 1 and 2 | 64 | LB | L | 969 | 1.000 |
| Cohort 1 | 32 | LB | LA | 231 | 1.000 |
| Cohort 2 | 32 | LB | LA | 242 | 1.000 |
| Cohorts 1 and 2 | 64 | LB | LA | 928 | 1.000 |
| Cohort 1 | 32 | LB | AL | 115 | 0.253 |
| Cohort 2 | 32 | LB | AL | 163 | 0.983 |
| Cohorts 1 and 2 | 64 | LB | AL | 536 | 0.048* |
| Cohort 1 | 32 | L | LA | 232 | 1.000 |
| Cohort 2 | 32 | L | LA | 223 | 1.000 |
| Cohorts 1 and 2 | 64 | L | LA | 924 | 1.000 |
| Cohort 1 | 32 | L | AL | 105 | 0.137 |
| Cohort 2 | 32 | L | AL | 165 | 0.988 |
| Cohorts 1 and 2 | 64 | L | AL | 519 | 0.032* |
| Cohort 1 | 32 | LA | AL | 144 | 0.799 |
| Cohort 2 | 32 | LA | AL | 193 | 1.000 |
| Cohorts 1 and 2 | 64 | LA | AL | 667 | 0.567 |

A, ahead (0°); AR, ahead-right (-30°); RA (-60°), right-ahead; R: right (-90°); RB, right-behind (-120°); BR, behind-right (-150°); B, behind (±180°); BL, behind-left (150°); LB, left-behind (120°); L, left (90°); LA, left-ahead (60°); AL, ahead-left (30°). $^{*}p_{corr.} < 0.05$; $^{**}p_{corr.} < 0.01$; $^{***}p_{corr.} < 0.001$.

Table S5: **Results of Wilcoxon signed rank test with Sidak correction for multiple comparisons to examine the relationship between allocentric starting orientation and allocentric memory performance.**

| **Cohort** | $n$ | **Orientation 1** | **Orientation 2** | $W$ | $p_{corr.}$ |
|---|---|---|---|---|---|
| Cohort 1 | 32 | N | NE | 214 | 1.000 |
| Cohort 2 | 32 | N | NE | 147 | 0.548 |
| Cohorts 1 and 2 | 64 | N | NE | 712 | 0.552 |
| Cohort 1 | 32 | N | E | 175 | 0.945 |
| Cohort 2 | 32 | N | E | 147 | 0.548 |
| Cohorts 1 and 2 | 64 | N | E | 614 | 0.116 |
| Cohort 1 | 32 | N | SE | 136 | 0.357 |
| Cohort 2 | 32 | N | SE | 60 | 0.001** |
| Cohorts 1 and 2 | 64 | N | SE | 379 | <0.001*** |
| Cohort 1 | 32 | N | S | 99 | 0.040* |
| Cohort 2 | 32 | N | S | 78 | 0.007* |
| Cohorts 1 and 2 | 64 | N | S | 328 | <0.001*** |
| Cohort 1 | 32 | N | SW | 104 | 0.057 |
| Cohort 2 | 32 | N | SW | 8 | <0.001*** |
| Cohorts 1 and 2 | 64 | N | SW | 215 | <0.001*** |
| Cohort 1 | 32 | N | W | 142 | 0.458 |
| Cohort 2 | 32 | N | W | 172 | 0.922 |
| Cohorts 1 and 2 | 64 | N | W | 613 | 0.114 |
| Cohort 1 | 32 | N | NW | 225 | 1.000 |
| Cohort 2 | 32 | N | NW | 254 | 1.000 |
| Cohorts 1 and 2 | 64 | N | NW | 940 | 1.000 |
| Cohort 1 | 32 | NE | E | 179 | 0.967 |
| Cohort 2 | 32 | NE | E | 263 | 1.000 |
| Cohorts 1 and 2 | 64 | NE | E | 884 | 1.000 |
| Cohort 1 | 32 | NE | SE | 130 | 0.269 |
| Cohort 2 | 32 | NE | SE | 165 | 0.848 |
| Cohorts 1 and 2 | 64 | NE | SE | 581 | 0.058 |
| Cohort 1 | 32 | NE | S | 97 | 0.034* |
| Cohort 2 | 32 | NE | S | 184 | 0.985 |
| Cohorts 1 and 2 | 64 | NE | S | 511 | 0.011* |
| Cohort 1 | 32 | NE | SW | 106 | 0.065 |
| Cohort 2 | 32 | NE | SW | 130 | 0.269 |
| Cohorts 1 and 2 | 64 | NE | SW | 461 | 0.003** |
| Cohort 1 | 32 | NE | W | 118 | 0.140 |
| Cohort 2 | 32 | NE | W | 248 | 1.000 |
| Cohorts 1 and 2 | 64 | NE | W | 791 | 0.941 |
| Cohort 1 | 32 | NE | NW | 258 | 1.000 |
| Cohort 2 | 32 | NE | NW | 171 | 0.914 |
| Cohorts 1 and 2 | 64 | NE | NW | 865 | 1.000 |
| Cohort 1 | 32 | E | SE | 201 | 1.000 |
| Cohort 2 | 32 | E | SE | 170 | 0.904 |
| Cohorts 1 and 2 | 64 | E | SE | 736 | 0.700 |
| Cohort 1 | 32 | E | S | 137 | 0.373 |
| Cohort 2 | 32 | E | S | 213 | 1.000 |
| Cohorts 1 and 2 | 64 | E | S | 692 | 0.431 |
| Cohort 1 | 32 | E | SW | 193 | 0.997 |
| Cohort 2 | 32 | E | SW | 164 | 0.835 |
| Cohorts 1 and 2 | 64 | E | SW | 691 | 0.426 |
| Cohort 1 | 32 | E | W | 197 | 0.999 |
| Cohort 2 | 32 | E | W | 208 | 1.000 |
| Cohorts 1 and 2 | 64 | E | W | 1024 | 1.000 |

*Continued on next page*

| **Cohort** | $n$ | **Orientation 1** | **Orientation 2** | $W$ | $p_{corr.}$ |
|---|---|---|---|---|---|
| Cohort 1 | 32 | E | NW | 186 | 0.989 |
| Cohort 2 | 32 | E | NW | 124 | 0.197 |
| Cohorts 1 and 2 | 64 | E | NW | 604 | 0.095 |
| Cohort 1 | 32 | SE | S | 209 | 1.000 |
| Cohort 2 | 32 | SE | S | 253 | 1.000 |
| Cohorts 1 and 2 | 64 | SE | S | 903 | 1.000 |
| Cohort 1 | 32 | SE | SW | 232 | 1.000 |
| Cohort 2 | 32 | SE | SW | 177 | 0.957 |
| Cohorts 1 and 2 | 64 | SE | SW | 804 | 0.967 |
| Cohort 1 | 32 | SE | W | 263 | 1.000 |
| Cohort 2 | 32 | SE | W | 153 | 0.658 |
| Cohorts 1 and 2 | 64 | SE | W | 834 | 0.994 |
| Cohort 1 | 32 | SE | NW | 169 | 0.894 |
| Cohort 2 | 32 | SE | NW | 84 | 0.012* |
| Cohorts 1 and 2 | 64 | SE | NW | 497 | 0.008** |
| Cohort 1 | 32 | S | SW | 247 | 1.000 |
| Cohort 2 | 32 | S | SW | 225 | 1.000 |
| Cohorts 1 and 2 | 64 | S | SW | 990 | 1.000 |
| Cohort 1 | 32 | S | W | 193 | 0.997 |
| Cohort 2 | 32 | S | W | 136 | 0.357 |
| Cohorts 1 and 2 | 64 | S | W | 656 | 0.250 |
| Cohort 1 | 32 | S | NW | 100 | 0.042* |
| Cohort 2 | 32 | S | NW | 74 | 0.005** |
| Cohorts 1 and 2 | 64 | S | NW | 332 | < 0.001*** |
| Cohort 1 | 32 | SW | W | 257 | 1.000 |
| Cohort 2 | 32 | SW | W | 134 | 0.326 |
| Cohorts 1 and 2 | 64 | SW | W | 753 | 0.795 |
| Cohort 1 | 32 | SW | NW | 113 | 0.103 |
| Cohort 2 | 32 | SW | NW | 61 | 0.001** |
| Cohorts 1 and 2 | 64 | SW | NW | 343 | < 0.001*** |
| Cohort 1 | 32 | NW | W | 154 | 0.676 |
| Cohort 2 | 32 | NW | W | 194 | 0.998 |
| Cohorts 1 and 2 | 64 | NW | W | 681 | 0.367 |

N, north; NE, north-east; E, east; SE, south-east; S, south; SW, south-west; W, west; NW, north-west.
*$p_{corr.} < 0.05$; **$p_{corr.} < 0.01$; ***$p_{corr.} < 0.001$.

Table S6: **Results of Wilcoxon signed rank test with Sidak correction for multiple comparisons to examine the relationship between allocentric starting orientation and egocentric memory performance.**

| **Cohort** | $n$ | **Orientation 1** | **Orientation 2** | $W$ | $p_{corr.}$ |
|---|---|---|---|---|---|
| Cohort 1 | 32 | N | NE | 131 | 0.283 |
| Cohort 2 | 32 | N | NE | 71 | 0.004* |
| Cohorts 1 and 2 | 64 | N | NE | 392 | <0.001*** |
| Cohort 1 | 32 | N | E | 157 | 0.728 |
| Cohort 2 | 32 | N | E | 67 | 0.003* |
| Cohorts 1 and 2 | 64 | N | E | 441 | 0.002* |
| Cohort 1 | 32 | N | SE | 68 | 0.003* |
| Cohort 2 | 32 | N | SE | 20 | <0.001*** |
| Cohorts 1 and 2 | 64 | N | SE | 185 | <0.001*** |
| Cohort 1 | 32 | N | S | 116 | 0.124 |
| Cohort 2 | 32 | N | S | 132 | 0.297 |
| Cohorts 1 and 2 | 64 | N | S | 493 | 0.007* |
| Cohort 1 | 32 | N | SW | 53 | <0.001*** |
| Cohort 2 | 32 | N | SW | 45 | <0.001*** |
| Cohorts 1 and 2 | 64 | N | SW | 182 | <0.001*** |
| Cohort 1 | 32 | N | W | 196 | 0.999 |
| Cohort 2 | 32 | N | W | 89 | 0.018* |
| Cohorts 1 and 2 | 64 | N | W | 563 | 0.039* |
| Cohort 1 | 32 | N | NW | 137 | 0.373 |
| Cohort 2 | 32 | N | NW | 40 | <0.001*** |
| Cohorts 1 and 2 | 64 | N | NW | 313 | <0.001*** |
| Cohort 1 | 32 | NE | E | 260 | 1.000 |
| Cohort 2 | 32 | NE | E | 234 | 1.000 |
| Cohorts 1 and 2 | 64 | NE | E | 988 | 1.000 |
| Cohort 1 | 32 | NE | SE | 126 | 0.219 |
| Cohort 2 | 32 | NE | SE | 192 | 0.997 |
| Cohorts 1 and 2 | 64 | NE | SE | 628 | 0.152 |
| Cohort 1 | 32 | NE | S | 207 | 1.000 |
| Cohort 2 | 32 | NE | S | 248 | 1.000 |
| Cohorts 1 and 2 | 64 | NE | S | 957 | 1.000 |
| Cohort 1 | 32 | NE | SW | 139 | 0.406 |
| Cohort 2 | 32 | NE | SW | 138 | 0.389 |
| Cohorts 1 and 2 | 64 | NE | SW | 552 | 0.030* |
| Cohort 1 | 32 | NE | W | 235 | 1.000 |
| Cohort 2 | 32 | NE | W | 210 | 1.000 |
| Cohorts 1 and 2 | 64 | NE | W | 895 | 1.000 |
| Cohort 1 | 32 | NE | NW | 264 | 1.000 |
| Cohort 2 | 32 | NE | NW | 196 | 0.999 |
| Cohorts 1 and 2 | 64 | NE | NW | 898 | 1.000 |
| Cohort 1 | 32 | E | SE | 157 | 0.728 |
| Cohort 2 | 32 | E | SE | 193 | 0.997 |
| Cohorts 1 and 2 | 64 | E | SE | 702 | 0.491 |
| Cohort 1 | 32 | E | S | 226 | 1.000 |
| Cohort 2 | 32 | E | S | 216 | 1.000 |
| Cohorts 1 and 2 | 64 | E | S | 1018 | 1.000 |
| Cohort 1 | 32 | E | SW | 127 | 0.231 |
| Cohort 2 | 32 | E | SW | 182 | 0.979 |
| Cohorts 1 and 2 | 64 | E | SW | 614 | 0.116 |
| Cohort 1 | 32 | E | W | 251 | 1.000 |
| Cohort 2 | 32 | E | W | 201 | 1.000 |
| Cohorts 1 and 2 | 64 | E | W | 902 | 1.000 |

*Continued on next page*

| **Cohort** | $n$ | **Orientation 1** | **Orientation 2** | $W$ | $p_{corr.}$ |
|---|---|---|---|---|---|
| Cohort 1 | 32 | E | NW | 237 | 1.000 |
| Cohort 2 | 32 | E | NW | 250 | 1.000 |
| Cohorts 1 and 2 | 64 | E | NW | 964 | 1.000 |
| Cohort 1 | 32 | SE | S | 198 | 0.999 |
| Cohort 2 | 32 | SE | S | 155 | 0.693 |
| Cohorts 1 and 2 | 64 | SE | S | 719 | 0.596 |
| Cohort 1 | 32 | SE | SW | 242 | 1.000 |
| Cohort 2 | 32 | SE | SW | 256 | 1.000 |
| Cohorts 1 and 2 | 64 | SE | SW | 1014 | 1.000 |
| Cohort 1 | 32 | SE | W | 172 | 0.922 |
| Cohort 2 | 32 | SE | W | 183 | 0.982 |
| Cohorts 1 and 2 | 64 | SE | W | 696 | 0.455 |
| Cohort 1 | 32 | SE | NW | 152 | 0.640 |
| Cohort 2 | 32 | SE | NW | 209 | 1.000 |
| Cohorts 1 and 2 | 64 | SE | NW | 710 | 0.540 |
| Cohort 1 | 32 | S | SW | 193 | 0.997 |
| Cohort 2 | 32 | S | SW | 136 | 0.357 |
| Cohorts 1 and 2 | 64 | S | SW | 633 | 0.167 |
| Cohort 1 | 32 | S | W | 210 | 1.000 |
| Cohort 2 | 32 | S | W | 262 | 1.000 |
| Cohorts 1 and 2 | 64 | S | W | 924 | 1.000 |
| Cohort 1 | 32 | S | NW | 205 | 1.000 |
| Cohort 2 | 32 | S | NW | 205 | 1.000 |
| Cohorts 1 and 2 | 64 | S | NW | 1015 | 1.000 |
| Cohort 1 | 32 | SW | W | 144 | 0.494 |
| Cohort 2 | 32 | SW | W | 144 | 0.494 |
| Cohorts 1 and 2 | 64 | SW | W | 554 | 0.032* |
| Cohort 1 | 32 | SW | NW | 122 | 0.176 |
| Cohort 2 | 32 | SW | NW | 185 | 0.987 |
| Cohorts 1 and 2 | 64 | SW | NW | 601 | 0.089 |
| Cohort 1 | 32 | NW | W | 240 | 1.000 |
| Cohort 2 | 32 | NW | W | 164 | 0.835 |
| Cohorts 1 and 2 | 64 | NW | W | 795 | 0.950 |

N, north; NE, north-east; E, east; SE, south-east; S, south; SW, south-west; W, west; NW, north-west.
$^{*}p_{corr.} < 0.05$; $^{**}p_{corr.} < 0.01$; $^{***}p_{corr.} < 0.001$.
